# Phylogenetic comparative methods on phylogenetic networks with reticulations

**DOI:** 10.1101/194050

**Authors:** Paul Bastide, Claudia Solís-Lemus, Ricardo Kriebel, K. William Sparks, Cécile Ané

## Abstract

The goal of Phylogenetic Comparative Methods (PCMs) is to study the distribution of quantitative traits among related species. The observed traits are often seen as the result of a Brownian Motion (BM) along the branches of a phylogenetic tree. Reticulation events such as hybridization, gene flow or horizontal gene transfer, can substantially affect a species’ traits, but are not modeled by a tree. *Phylogenetic networks* have been designed to represent reticulate evolution. As they become available for downstream analyses, new models of trait evolution are needed, applicable to networks. One natural extension of the BM is to use a weighted average model for the trait of a hybrid, at a reticulation point. We develop here an efficient recursive algorithm to compute the phylogenetic variance matrix of a trait on a network, in only one preorder traversal of the network. We then extend the standard PCM tools to this new framework, including phylogenetic regression with covariates (or phylogenetic ANOVA), ancestral trait reconstruction, and Pagel’s λ test of phylogenetic signal. The trait of a hybrid is sometimes outside of the range of its two parents, for instance because of hybrid vigor or hybrid depression. These two phenomena are rather commonly observed in present-day hybrids. Transgressive evolution can be modeled as a shift in the trait value following a reticulation point. We develop a general framework to handle such shifts, and take advantage of the phylogenetic regression view of the problem to design statistical tests for ancestral transgressive evolution in the evolutionary history of a group of species. We study the power of these tests in several scenarios, and show that recent events have indeed the strongest impact on the trait distribution of present-day taxa. We apply those methods to a dataset of *Xiphophorus* fishes, to confirm and complete previous analysis in this group. All the methods developed here are available in the **Julia** package **PhyloNetworks.**

---

The evolutionary history of species is known to shape the present-day distribution of observed characters (Felsenstein 1985). Phylogenetic Comparative Methods (PCMs) have been developed to account for correlations induced by a shared history in the analysis of a quantitative dataset (Pennell and Harmon 2013). They usually rely on two main ingredients: a time-calibrated phylogenetic tree, and a dynamical model of trait evolution, that should be chosen to capture the features of the trait evolution over time. Much work has been made on the second ingredient, with more and more sophisticated models of trait evolution, with numerous variations around the original Brownian Motion (BM), see for instance Felsenstein (1985); Hansen and Martins (1996); Hansen (1997); Blomberg et al. (2003); Butler and King (2004); Beaulieu et al. (2012); Landis et al. (2013); Blomberg (2016), to cite only but a few.

In contrast, the first assumption has not been questioned until now (but see Jhwueng and O’Meara 2018). However, phylogenetic trees are not always well suited to capture relationships between species, and *phylogenetic networks* are sometimes needed. Phylogenetic networks differ from trees by added reticulation points, where two distinct branches come together to create a new species. Such reticulations can represent various biological mechanisms, like hybridization, gene flow or horizontal gene transfer, that are known to be common in some groups of organisms (Mallet 2005, 2007). Ignoring those events can lead to misleading tree inference (Kubatko 2009; Solís-Lemus et al. 2016; Long and Kubatko 2018). Thanks to recent methodological developments, the statistical inference of reliable phylogenetic networks has become possible (Maddison 1997; Degnan and Salter 2005; Kubatko 2009; Yu et al. 2012, 2014; Yu and Nakhleh 2015; Solís-Lemus and Ané 2016). Although these state-of-the-art methods are still limited by their computational burden, we believe that the use of these networks will increase in the future. The goal of this work is to propose an adaptation of standard PCMs to a group of species with reticulate evolution, related by a network instead of a tree.

We describe an extension of the BM model of trait evolution to a network. The main modeling choice is about the trait of hybrid species. How should these species inherit their trait from their two parents? In this work, we first choose a weighted-average merging rule: the trait of a hybrid is a mixture of its two parents, weighted by their relative genetic contributions. This rule can be seen as a reasonable null model. However, in some cases, the trait of a hybrid is observed to be outside of the range of its two parents. This phenomenon can be modeled by a *shift* in the trait value occurring right after the reticulation point: the hybrid trait value being the weighted average of the two parents, plus an extra term specific to the hybridization event at hand. Such a shift can model several biological mechanisms, such as transgressive segregation (Rieseberg et al. 1999) or heterosis (Fiévet et al. 2010; Chen 2013), with hybrid vigor (when the hybrid species is particularly fit to its environment) or depression (when the hybrid is ill-fit). In the following, we will refer to this class of phenomena using the generic term transgressive evolution. Here, this term only refers to the hybrid trait being different from the weighted average of its parents. This model allows for an explicit mathematical derivation of the trait distribution at the tips of the network and extends to networks the use of standard PCM tools such as phylogenetic regression (Grafen 1989, 1992), ancestral state reconstruction (Felsenstein 1985; Schluter et al. 1997) or tests of phylogenetic signal (Pagel 1999).

In the following, we first describe this BM model of trait evolution and show how it fits into the standard PCM framework. We then show how to add shifts in the trait values to model transgressive evolution. We propose a statistical test for transgressive evolution. These methods are validated with a simulation study, and with the theoretical study of the power of the tests in a range of scenarios. Finally, we revisit the analysis of a *Xiphophorus* dataset about sword index and female preference made by Cui et al. (2013), in the light of our new network methods.

## Model

In our model for trait evolution on a phylogenetic network, the novel aspect is the merging rule at reticulation events, compared to standard PCMs on trees. Our model is very similar to that defined in Jhwueng and O’Meara (2018), but we adopt a different statistical view point, based on the phylogenetic linear regression representation.

### Trait Evolution on Networks

#### Phylogenetic Network

In this work, we assume that we have access to a *rooted, calibrated and weighted phylogenetic network* that describes the relationships between a set of observed species (Huson et al. 2010). In such a network, reticulations, or *hybrids*, are nodes that have two parent nodes. They receive a given proportion of their genetic material from each parent. This proportion is controlled by a weight *γ_e_* that represents the *inheritance probability* associated with each branch *e* of the network. A branch that is *tree-like*, i.e. that ends at a non-hybrid node, has a weight *γ_e_* = 1. We further assume that the length *ℓ_e_* of a branch *e* represents evolutionary time. In the example in Figure 1a, the two hybrid edges have length zero, but this need not to be the case, see Jhwueng and O’Meara (2018); Degnan (2017).

**Figure 1:**
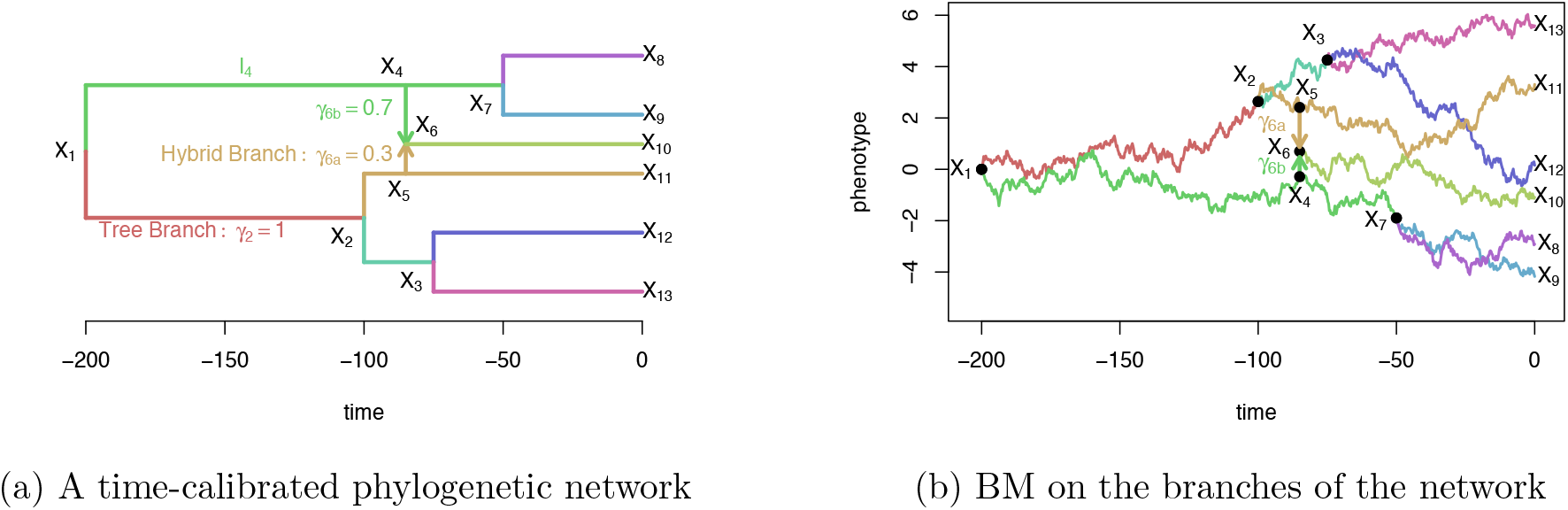
Realization of a BM (with *µ* = 0 and *σ*^2^ = 0.04) on a calibrated network. Only tip values are observed (here at time *t* = 0). For simplicity, the two hybrid branches were chosen to have a length of 0, but this need not be the case. Inheritance probabilities at the hybridization event are *γ*_6*a*_ and *γ*_6*b*_, with *γ*_6*a*_ + *γ*_6*b*_ = 1.

#### Brownian Motion Model

Since Felsenstein (1985), the Brownian Motion (BM) has been intensively used to model trait evolution on phylogenetic trees. It is well adapted to model several biological processes, from random genetic drift, to rapid adaptation to a fluctuating environment (see e.g. Felsenstein 2004, Chap. 24). In order to adapt this process to a network instead of a tree, we define a weighted average merging rule at hybrids, as defined below. This rule expresses the idea that a hybrid inherits its trait from both its parents, with a relative weight determined by the proportion of genetic material received from each: if it inherits 90% of its genes from parent *A*, then 90% of its trait value should be determined by the trait of *A*. Because the BM usually models the evolution of a polygenic character, that is the additive result of the expression of numerous genes, this rule is a natural null hypothesis.

##### Definition 1

(BM on a Network). Consider a rooted phylogenetic network with branch lengths and inheritance probabilities. Let *X_v_* be the random variable describing the trait value of node (or vertex) *v*.

- At the root node *ρ*, we assume that *X_ρ_* = *µ* is fixed.
- For a tree node *v* with parent node *a*, we assume that *X_v_* is normally distributed with mean *X_a_* + ∆_*e*_ and with variance *σ*^2^*ℓ_e_*, with *σ*^2^ the variance rate of the BM, and *ℓ_e_* the length of the parent edge *e* from *a* to *v*. ∆_*e*_ is a (fixed) shift value associated with branch *e*, possibly equal to 0.
- For a hybrid node *v* with parent nodes *a* and *b*, we assume that *X_v_* is normally distributed with mean *γ_e_a__ X_a_* + *γ_e_b__ X_b_*, where *e_a_* and *e_b_* are the hybrid edges from *a* (and *b*) to *v*. If these edges have length 0, meaning that *a*, *b* and their hybrid *v* are contemporary, then *X_v_* is assumed to have variance 0, conditional on the parent traits *X_a_* and *X_b_*. In general, the conditional variance of *X_v_* is *γ_e_a__ σ^2^ ℓ_e_a__* + *γ_e_b__ σ ℓ_e_b__*. For the sake of identifiability, shifts are not allowed on hybrid branches (see Section on Transgressive Evolution for further details).

An example of such a process (without shift) is presented Figure 1b. This process is similar to Jhwueng and O’Meara (2018), except that the shifts are treated differently. See Section on Transgressive Evolution and Discussion for more information on the links and differences between the two models. For the sake of generality, shifts are allowed on any tree edge. We will see in the next section how they are used to model transgressive evolution. In the rest of this section, we take all shifts to be zero, and only consider the un-shifted BM (∆_*e*_ = 0 for all edges *e*).

Note that the state at the root, *µ*, could also be drawn from a Gaussian distribution, instead of being fixed. This would not change the derivations below, and would simply add a constant value to all terms in the variance matrix.

### Variance Matrix

#### From a Tree to a Network

The distribution of trait values at all nodes, **X**, can be fully characterized as a multivariate Gaussian with mean *µ***1**_*m*+*n*_ and variance matrix *σ*^2^**C**, where **1**_*m*+*n*_ is the vector of ones, *n* is the number of tips and *m* the number of internal nodes. The matrix **C**, which depends on the topology of the network, encodes the correlations induced by the phylogenetic relationships between taxa. When the network reduces to a tree (if there are no hybrids), then **C** is the well-known BM covariance (Felsenstein 1985): **C**_*ij*_ = *t_ij_* is the time of shared evolution between nodes *i* and *j*, i.e. the time elapsed between the root and the most recent common ancestor (MRCA) of *i* and *j*.

When the network contains hybrids, this formula is not valid anymore. To see this, let’s re-write *t_ij_* as:

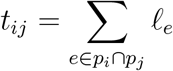

where *p_i_* is the path going from the root to node *i*. This formula just literally expresses that *t_ij_* is the length of the shared path between the two nodes, that ends at their MRCA. On a network, the difficulty is that there is *not a unique path* going from the root to a given node. Indeed, if there is a hybrid among the ancestors of node *i*, then a path might go “right” of “left” of the hybrid loop to go from the root to *i*.

Under the BM model in Definition 1 (with a fixed root), it turns out that we need to *sum* over all the possible paths going from the root to a given node, weighting paths by the inheritance probabilities *γ_e_* of all the traversed edges:

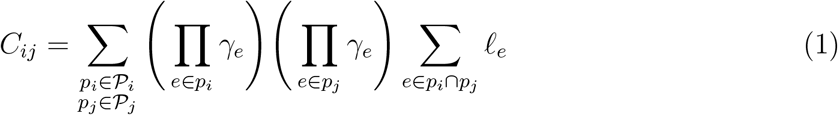

where *P_i_* denotes the set of all the paths going from the root to node *i*.

This general formula for **C** was first presented in Pickrell and Pritchard (2012) in the context of population genomics. A formal proof is provided here (Appendix).

*Remark* 1 (Variance reduction). From the expression above, we can show that the variance of any tip *i* decreases with each hybridization ancestral to *i*. Consider a time-consistent network, in the sense that *all* paths from the root to a given hybrid node have the same length, as expected if branch lengths measure calendar time. Note that this is the opposite of the “NELP” property (No Equally Long Paths) defined by Pardi and Scornavacca (2015). For tip *i*, let *t_i_* be the length of any path from the root to *i*. If the network is a tree, then *C_ii_* = *t_i_*. If the history of tip *i* involves one or more reticulations, then we show (Appendix), that:

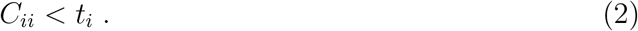

This shows that hybridization events, which imply taking a weighted means of two traits, cause the trait variance to decrease. Note that this variance reduction is a consequence of our particular model of trait hybridization. Other merging rules might yield different trait variances after hybrid nodes. Our model of transgressive evolution acts on the trait mean (through shifts ∆_*e*_, see next section) such that variation due to transgressive segregation is assumed to be captured by variation in the trait means, not by an increased trait variance.

#### Algorithm

The formula above, although general, is not practical to compute. Using the recursive characterization of the process given in Definition 1, we can derive an efficient way to compute this covariance matrix across all nodes in the network (tips and internal nodes), in a single traversal of the network. This traversal needs to be in “preorder”, from the root to the tips, such that any given node is listed after all of its parent(s): for any two nodes numbered *i* and *j*, if there is a directed path from *i* to *j*, then *i* ≤ *j*. Such an ordering (also called topological sorting) can be obtained in linear time in the number of nodes and edges (Kahn 1962). On Figure 1a, nodes are numbered from 1 to 13 in preorder. The result below, proved in the Appendix, provides an efficient algorithm to compute the phylogenetic variance matrix **C** in a time linear in the number of nodes of the network, in a single preorder traversal.

##### Proposition 1

(Iterative computation of the phylogenetic variance). *Assume that the nodes of a network are numbered in preorder. Then* **C** *can be calculated using the following step for each node i, from i* = 1 *to i* = *n* + *m:*

- *If i* = 1 *then i is the root, and C_ii_* = 0.
- *If i is a tree node, denote by a the parent of i, and by ℓ_e_a__ the length of the branch e_a_ going from a to i. Then:*

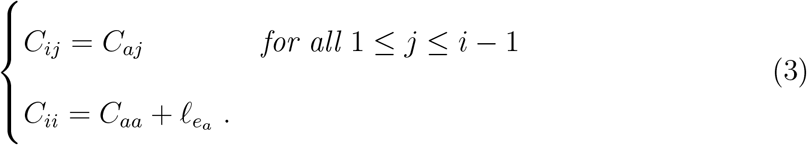
- *If i is a hybrid node, denote by a and b the parents of i, by ℓ_e_a__ and ℓ_e_b__ the lengths of the branches e_a_ and e_b_ going from a or b to i, and by γ_e_a__ and γ_e_b__ the associated inheritances probabilities. Then:*

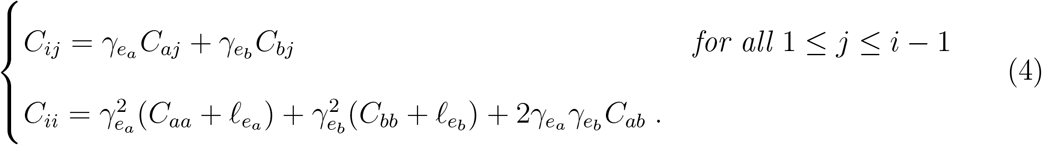

### Phylogenetic Regression

We can now define a *phylogenetic regression* on networks, the same way it is defined for phylogenetic trees (Grafen 1989, 1992).

#### Linear Regression Framework

Define **Y** as the vector of trait values observed at the tips of the network. This is a sub-vector of the larger vector of trait values at all nodes. Let **C**^tip^ be the sub-matrix of **C**, with covariances between the observed taxa (tips). The phylogenetic linear regression can be written as:

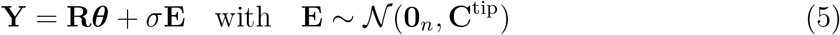

where **R** is a *n × q* matrix of regressors, and ***θ*** a vector of *q* coefficients. We can recover the distribution of **Y** under a simple BM with a fixed root value equal to *µ* (and no shift) by taking **R**=**1**_*n*_ and ***θ***=*µ* (with *q* = 1). The regression matrix **R** can also contain some explanatory trait variables of interest. In this phylogenetic regression, the BM model applies to the residual variation not explained by predictors, **E**.

This formulation is very powerful, as it recasts the problem into the well-known linear regression framework. The variance matrix **C**^tip^ is known (it is entirely characterized by the network used) so that, through a Cholesky factorization, we can reduce this regression to the canonical case of independent sampling units. This problem hence inherits all the features of the standard linear regression, such as confidence intervals for coefficients or data prediction, as explained in the next paragraph.

#### Ancestral State Reconstruction and Missing Data

The phylogenetic variance matrix can also be used to do ancestral state reconstruction, or missing data imputation. Both tasks are equivalent from a mathematical point of view, rely on the Best Linear Unbiased Predictor (BLUP, see e.g. Robinson 1991) and are well known in the standard PCM toolbox. They have been implemented in many **R** packages, such as **ape** (Paradis et al. 2004, function ace), **phytools** (Revell 2012, function **fastAnc**) or **Rphylopars** (Goolsby et al. 2017, function **phylopars**). In our **Julia** package **PhyloNetworks**, it is available as function **ancestralStateReconstruction**.

#### Pagel’s λ

The variance structure induced by the BM can be made more flexible using standard transformations of the network branch lengths, such as Pagel’s *λ* (Pagel 1999). Because the network is calibrated with node ages, it is time-consistent: the time *t_i_* elapsed between the root and a given node *i* is well defined, and does not depend on the path taken. Hence, the lambda transform used on a tree can be extended to networks, as shown below.

##### Definition 2

(Pagel’s *λ* transform). First, for any hybrid tip in the network, add a child edge of length 0 to change this tip into an internal (hybrid) node, and transfer the data from the former hybrid tip to the new tip. Next, let *e* be a branch of the network, with child node *i*, parent node pa(*i*), and length *ℓ_e_*. Then its transformed length is given by:

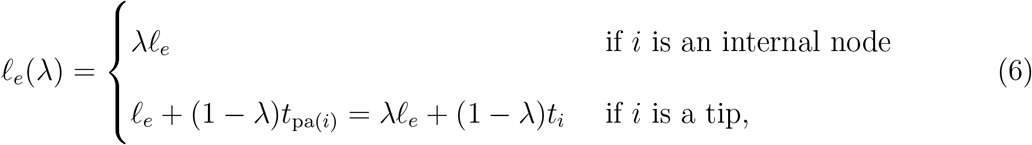

where *t_i_* and *t*_pa(*i*)_ are the times elapsed from the root to node *i* and to its parent.

The interpretation of this transformation in term of phylogenetic signal is as usual: when *λ* decreases to zero, the phylogenetic structure is less and less important, and traits at the tips are completely independent for *λ* = 0. The first step of resolving hybrid tips is similar to a common technique to resolve polytomies in trees, using extra branches of length 0. This transformation does not change the interpretation of the network or the age of the hybrid. The added external edge does allow extra variation specific to the hybrid species, however, immediately after the hybridization, by Pagel’s *λ* transformation. The second part of (6) applies to the new external tree edge, and hybrid edges are only affected by the first part of (6). The transformation’s impact on the matrix **C**^tip^ is not exactly the same as on trees. It still involves a simple multiplication of the off-diagonal terms by *λ*, but the diagonal terms are also modified. The following proposition is proved in the Appendix.

##### Proposition 2

(Pagel’s *λ* effect on the variance). *The phylogenetic variance of a BM running on a network transformed by a parameter λ*, **C**(*λ*) *is given by:*

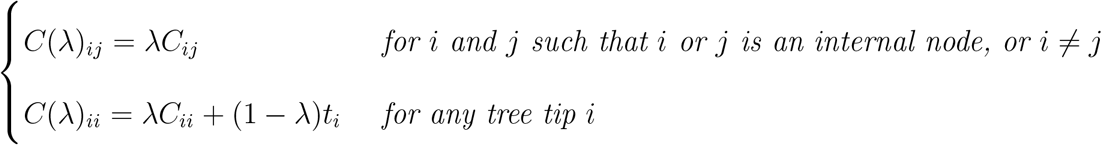

*where* **C** = **C**(1) *is the variance of the BM process on the non-transformed network*.

On a tree, we have *C*(*λ*)_*ii*_ = *t_i_* for any tip *i* and any *λ*, so that the diagonal terms remain unchanged. This is not true on a network, however, as the Pagel transformation erases the variance-reduction effect of ancestral hybridizations.

Other transformations, for instance based on Pagel’s *κ* or *δ* (Pagel 1999), could be adapted to the phylogenetic network setting. Although these are not implemented for the moment, they would be straightforward to add in our linear regression framework.

### Shifted BM and Transgressive Evolution

In our BM model, we allowed for shifts on non-hybrid edges. In this section, we show how those shifts can be inferred from the linear regression framework, and how they can be used to test for ancestral transgressive evolution events. When considering shifts, we again require that all tips are tree nodes. If a tip is a hybrid node (with two parents), then the network is first resolved by adding a child edge of length 0 to the hybrid, making this node an internal node. This network resolution does not affect the interpretation of the network or the variance of the BM model. It adds more flexibility to the mean vector of the BM process, because the extra edge is a tree edge on which a shift can be placed.

#### Shift Vector

We first describe an efficient way to represent the shifts on the network branches in a vector format. In Definition 1, we forbade shifts on hybrid branches. This does not lose generality, and is just for the sake of identifiability. Indeed, a hybrid node connects to three branches, two incoming (the hybrid edges) and one outgoing (a tree edge typically). A shift on any of these three branches would impact the same set of nodes (apart from the hybrid itself), and hence would produce the same data distribution at the tips. The position of a shift on these three branches is consequently not identifiable. By restricting shifts to tree branches, the combined effect of branches with the same set of descendants is identified by a shift on a single (tree) edge. We can combine all shift values in a vector **∆** indexed by nodes:

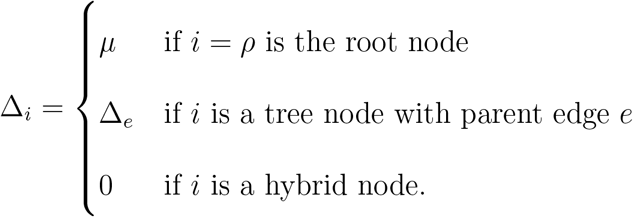

Note that any tree edge *e* is associated to its child node *i* in this definition. In the following, when there is no ambiguity, we will refer indifferently to one or the other.

#### Descendence Matrix

For a rooted tree, a matrix of 0/1 values where each column corresponds to a clade can fully represent the tree topology. In column *j*, entries are equal to 1 for descendants of node number *j*, and 0 otherwise (Ho and Ané 2014; Bastide et al. 2017). This representation is similar to the additive binary coding of a tree (Farris et al. 1970; Brooks 1981) as used for instance in methods by matrix representation parsimony for supertree estimation (Baum 1992; Ragan 1992) On a network, a node *i* can be a “partial” descendant of *j*, with the proportion of inherited genetic material represented by the inheritance probabilities *γ_e_*. Hence, the descendence matrix of a network can be defined with non-binary entries between 0 and 1 as follows.

##### Definition 3

(Descendence Matrix). The descendence matrix **U** of a network, given some ordering of its *n* tips and *m* internal nodes, is defined as an (*n* + *m*) *×* (*n* + *m*) matrix by:

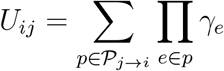

where *𝒫_j→i_* is the set of all the paths going from node *j* to node *i* (respecting the direction of edges). Note that, if *i* is not a descendant of *j*, then *𝒫_j→i_* is empty and *U_ij_* = 0. By convention, if *i* = *j*, we take *U_ii_* = 1 (that is, a node is considered to be a descendant of itself). If the network is a tree, we recover the usual definition (all the *γ_e_* are equal to 1).

In general, the set of nodes *i* for which *U_ij_ >* 0 is the hardwired cluster of *i*, or the clade below *i* if the network is a tree.

Further define **T** as the (non-square) submatrix of **U** made of the rows that correspond to tip nodes (see example below).

*Example* 1 (Descendence Matrix and Shift Vector). The descendence matrices **U** and **T** associated with the network presented in Figure 2 are shown below, with zeros replaced by dots to improve readability:

**Figure 2.**
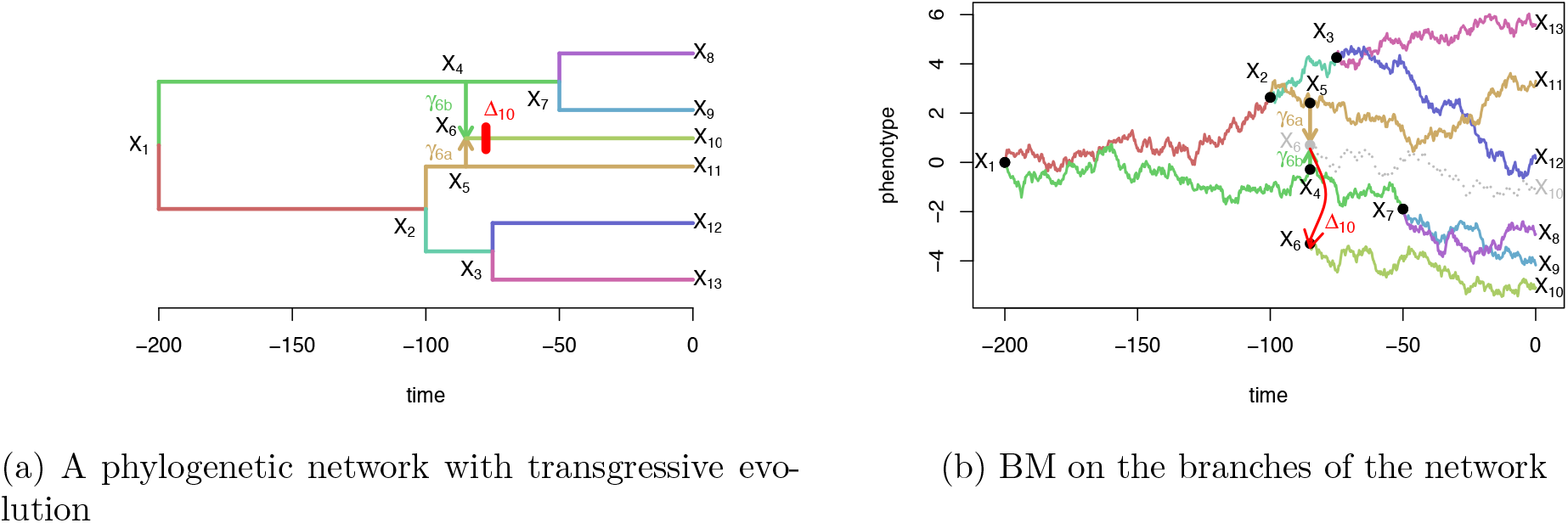
Realization of a univariate BM process (with *µ* = 0 and *σ*^2^ = 0.04) on a calibrated network, with transgressive evolution. The shift occurs right after the hybridization event, and changes the trajectory of the BM from the grey dotted one to the colored one.

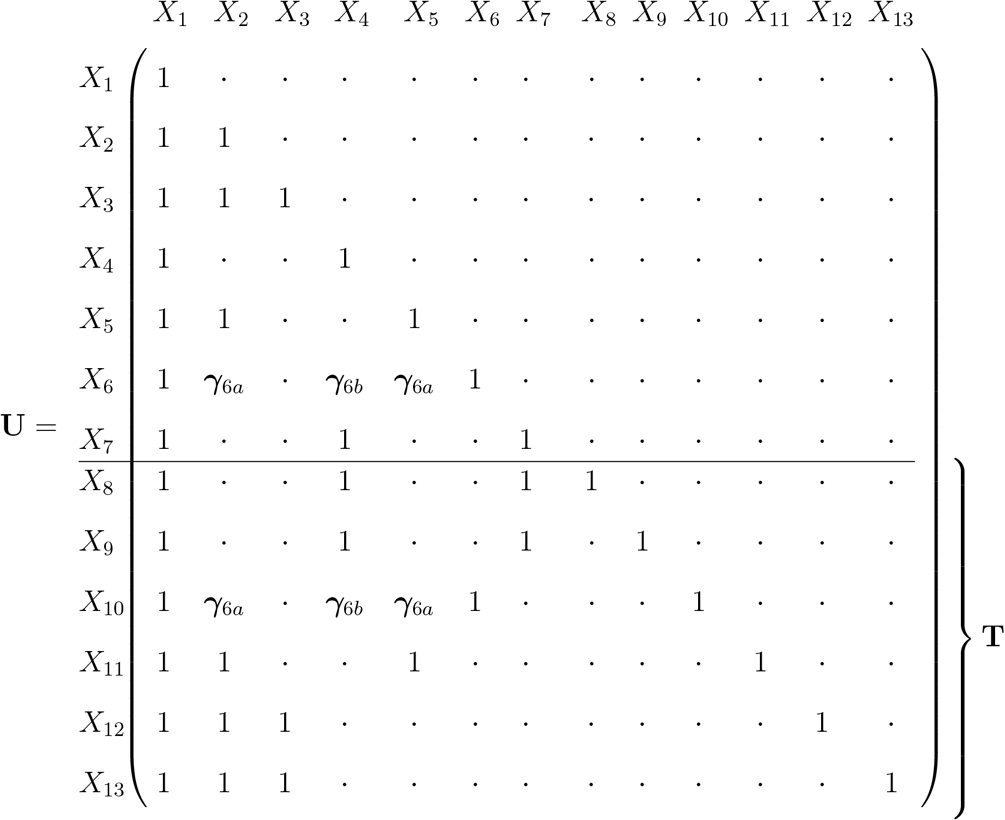

The associated shift vector and associated trait means at the tips are shown below, where the only non-zero shift is assumed to correspond to transgressive evolution at the hybridization event, captured by ∆_10_ on edge 10:

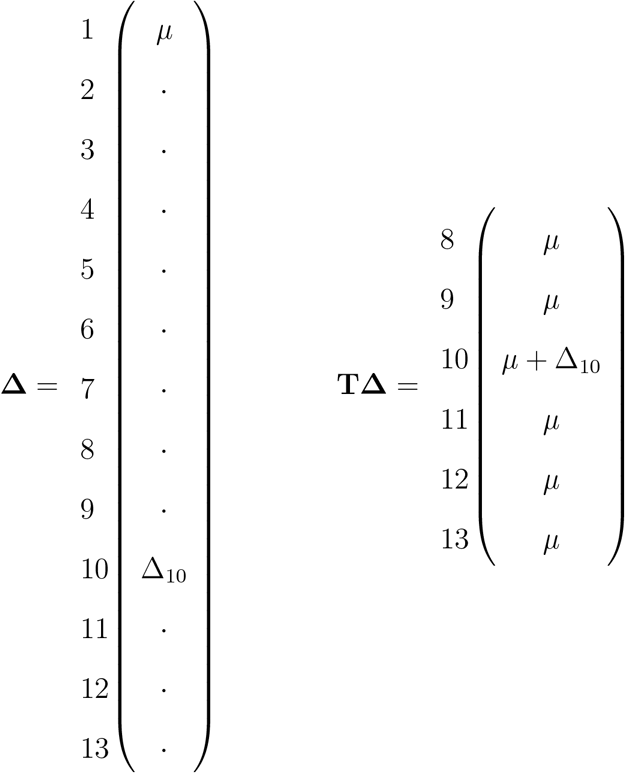

Note that rapid trait evolution or jumps in the trait value in other parts of the phylogeny could be also be modeled, by letting ∆_*i*_ be non-zero for other tree edges *i*. However, allowing for too many non-zero values in **∆** can lead to severe identifiability issues. See e.g. Bastide et al. (2017) for an identifiability study of this vector on a phylogenetic tree.

#### Linear Model

The shifted BM model in Definition 1 can be expressed by:

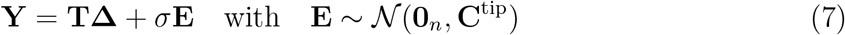

where **Y** is the trait vector at the tips, and **∆** and **T** are the shift vector and the descendence matrix as defined above (see the Appendix for the proof).

#### Transgressive Evolution

Even though the linear formulation above makes it easier to study, the problem of locating the non-zero shifts on the branches of a phylogenetic tree is difficult, and is still an active research area (see e.g. Uyeda and Harmon 2014; Bastide et al. 2017; Khabbazian et al. 2016; Bastide et al. 2018).

On networks as on trees, a shift can represent various biological processes. In the present work, we limit our study to shifts occurring on branches that are outgoing from a hybrid node (see Figure 2 for an example). Such shifts might represent a *transgressive evolution* effect, as defined in the introduction, and as a component of hybridization: the new species inherits its trait as a weighted average of the traits of its two parents, plus a shift representing extra variation, perhaps as a result of rapid selection.

Limiting shifts to being right after reticulations avoids the difficult exploration of all the possible locations of an unknown number of shifts on all the tree branches.

### Statistical Tests for Transgressive Evolution

As there are typically only a few hybridization events in a phylogenetic network, we can test for transgressive evolution on each one individually. Thanks to the linear framework described above, this amounts to a well-known test of fixed effects.

#### Statistical Model

Denote by **N** the *n × h* sub-matrix of **T** containing only the columns corresponding to tree branches outgoing from hybrid nodes. We assume that **N** has full rank, that is, that the transgressive evolution shifts are identifiable. This is likely to be the case in networks that can be inferred by current methods, which typically have a small number of reticulations. We further denote by **N̅** the vector of size *n* containing the row sums of **N**: for tip *i*, 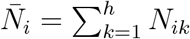. Then the phylogenetic linear regression extending (5) with transgressive evolution can be written as:

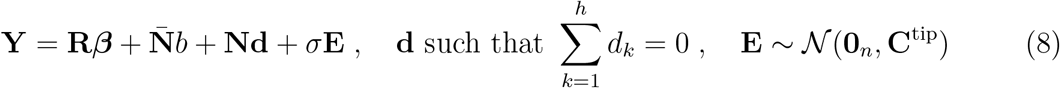

where **R** is a given matrix of regressors, with associated coefficients ***β***. These are included for the sake of generality, but usually only represent the ancestral state of the BM: **R** = **1**_*n*_ and ***β*** = *µ*. The coefficient *b* represents a *common* transgressive evolution effect, that would affect all the hybridization events uniformly, while the vector **d** has *h* entries with a specific deviation from this common effect for each event, and represents *heterogeneity*.

#### F Test

When written this way, the problem of testing for transgressive evolution just amounts to testing the fixed effects *b* and **d**. Some hypotheses that can be tested are summarized in the next table. *𝓗*_0_ corresponds to the null model where the hybrids inherit their parents weighted average. *𝓗*_1_ is a model where all hybridization events share the same transgressive evolution effect, the trait being shifted by a common coefficient *b*. Finally, *𝓗*_2_ is a model where each hybridization event *k* has its own transgressive evolution effect, with a shift *b* + *d_k_*.

|  | Hypotheses | Linear Model |
| --- | --- | --- |
| $\mathcal{H}_0$ | No transgressive evolution | $b = 0$ and $\mathbf{d} = \mathbf{0}_h$ |
| $\mathcal{H}_1$ | Single effect transgressive evolution | $b \neq 0$ and $\mathbf{d} = \mathbf{0}_h$ |
| $\mathcal{H}_2$ | Multi effect transgressive evolution | $b \neq 0$ and $\mathbf{d} \neq \mathbf{0}_h$ |

Tests of fixed effects are very classic in the statistics literature (see e.g. Lehman 1986; Searle 1987). Compared to a likelihood ratio test, an F-test is exact and is more powerful, when available. Here we can define two F statistics *F*_10_ and *F*_21_ (see the Appendix). To see if *𝓗*_2_ fits the data significantly better than *𝓗*_1_, we compare *F*_21_ to an F distribution with degrees of freedom *r*_[**R N**]_ *− r*_[**R N̅** ]_ and *n − r*_[**R N**]_, where *r* is the matrix rank, and [**R N**] is the matrix obtained by pasting the columns of **R** and **N** together. To test *𝓗*_1_ versus the null model *𝓗*_0_, we compare *F*_10_ to an F distribution with degrees of freedom *r*_[**R N̅** ]_ *− r*_**R**_ and *n − r*_[**R N̅**]_. We study these tests for several symmetric networks in the following section.

## Simulation and Power Study

In this section, we first analyse the performance of the PCM tools described above, and then provide a theoretical power study of our statistical tests for transgressive evolution.

### Implementation of the Network PCMs

All the tools described above, as well as simulation tools, were implemented in the Julia (Bezanson et al. 2017) package **PhyloNetworks** (Solís-Lemus et al. 2017). To perform a phylogenetic regression, the main function is **phyloNetworklm**. It relies on functions **preorder!** and **sharedPathMatrix** to efficiently compute the variance matrix using the algorithm in Proposition 1, and on Julia package **GLM** (Bates 2016) for the linear regression. All the analysis and extraction tools provided by this **GLM** package can hence be used, including the **ftest** function to perform the F statistical tests for transgressive evolution. For the *Xiphophorus* fishes study (see below), we developed a function **calibrateFromPairwiseDistances!** to calibrate a network topology based on pairwise genetic distances.

### Simulation Study

#### Setting

We considered 4 network topologies, all based on the same symmetric backbone tree with unit height and 32 tips, to which we added several hybridization events (Fig. 3, top). Those events were either taken very recent and numerous (*h* = 8 events each affecting 1 taxon) or quite ancient and scarce (*h* = 2 events each affecting 4 taxa). All networks had 8 tips with a hybrid ancestry. All the hybridization events had inheritance probability *γ* = 0.3. We then simulated datasets on these networks with *µ* = 0, *σ*^2^ = 1, and Pagel’s *λ* transformation with *λ* in {0, 0.25, 0.5, 0.75, 1}. Recall that *λ* = 0 corresponds to all tips being independent, and *λ* = 1 is the simple BM on the original network. Each simulation scenario was replicated 500 times. To study the scalability of the implementation, we then reproduced these analysis on networks with 32 to 256 tips, and 1 to 8 hybridization events, each affecting 8 tips.

**Figure 3.**
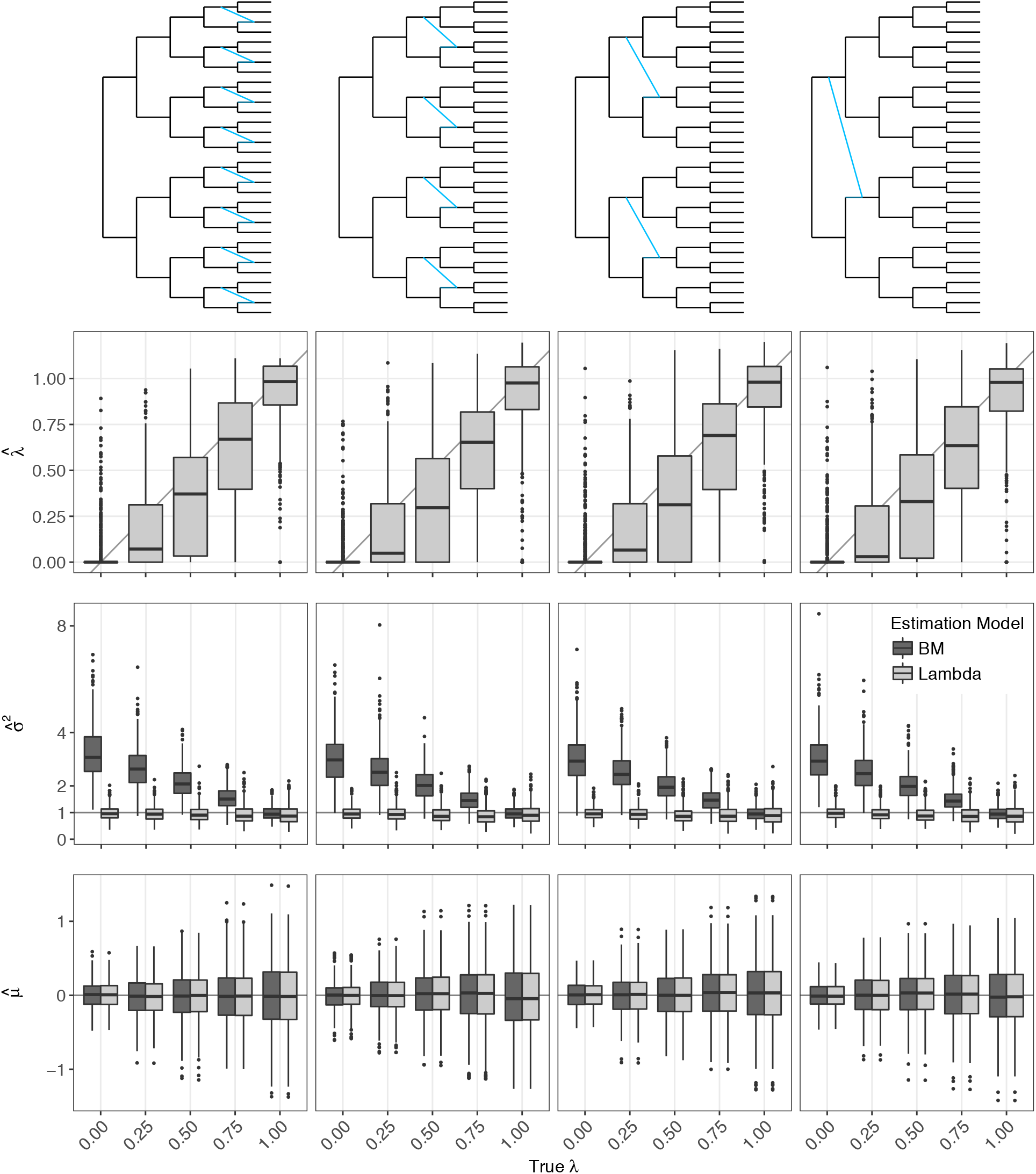
Estimated *λ*, *σ*^2^ and *µ* values for several network topologies, with *γ* = 0.3, when the data are simulated according to a BM process with Pagel’s *λ* transformation. Data were analyzed either with a straight BM model, which corresponds to *λ* = 1 (dark grey), or with Pagel’s *λ* transformed model (light grey). True values are marked by a grey line. Boxplots show variation across 500 replicates.

We analysed each dataset assuming either a BM or a *λ* model of evolution. When *λ*≠1, we could study the effect of wrongly using the BM. All the analyses were conducted on a laptop computer, with four Intel Core i7-6600U, and a 2.60GHz CPU speed.

#### Results

When the vanilla BM model is used for both the siWe considered 4 netwmulation and the inference, the two parameters *µ* and *σ*^2^ are well estimated, with no bias, for all the network topologies tested (Fig. 3, last two rows, dark grey boxes for true *λ* = 1). The estimation of *µ* is quite robust to the misspecification of the model, while *σ*^2^ tends to be over-estimated (Fig. 3, last two rows, dark grey boxes for true *λ* ≠ 1). This is expected, as in this case, the BM model wrongly tries to impose a strong correlation phylogenetic structure on the data, and can only account for the observed diversity by raising the estimated variance, to accommodate both phylogenetic variance and independent intra-specific variation. When we use the true *λ* model for the inference, this bias is corrected, and both *µ* and *σ*^2^ are correctly estimated (Fig. 3, last two rows, light grey boxes). Furthermore, the *λ* estimate has a small bias but rather high variance (Fig. 3, second row). As expected, when the number of taxa increases, this variance decreases (data not shown). Finally, our implementation is quite fast (Fig. 4), with computing times ranging between 1 and 10 ms for a BM fit, and between 10 ms and 1 s for a Pagel’s *λ* fit.

**Figure 4.**
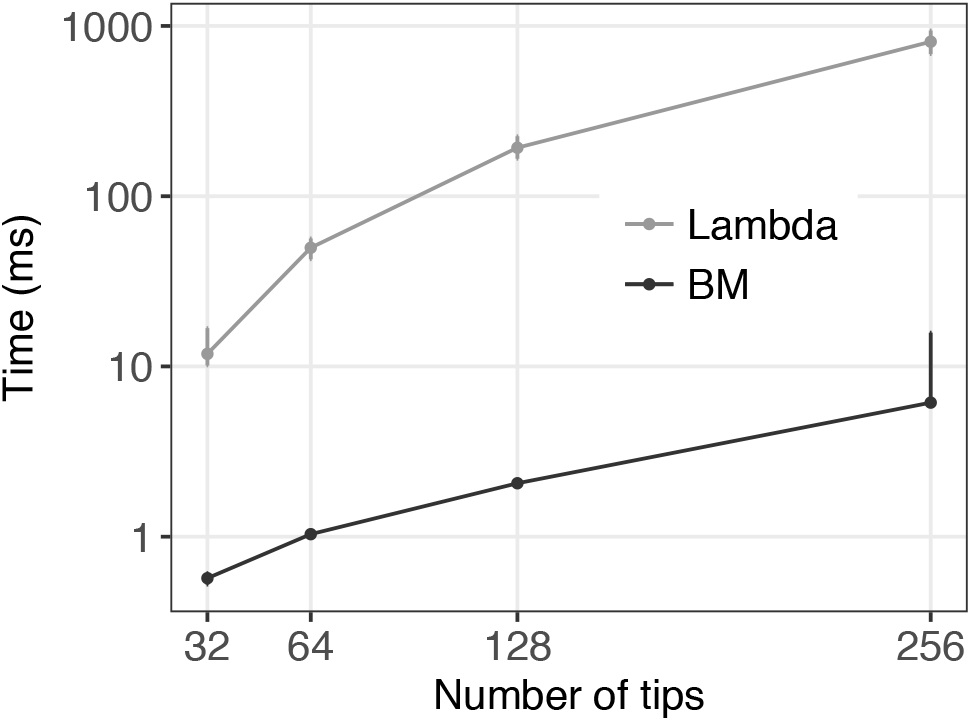
Computing time needed for fitting a continuous trait evolution model in **PhyloNetworks**. Median and confidence interval for 6000 repetitions in various conditions for each number of taxa. A log scale is used for the computing time.

### Power Study of the Statistical Tests for Transgressive Evolution

We determined that our test statistics have the following noncentral Fisher-Snedecor (F) distributions:

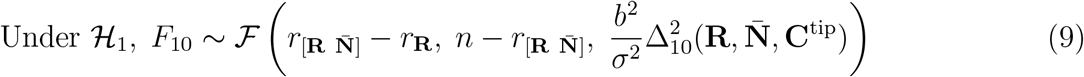

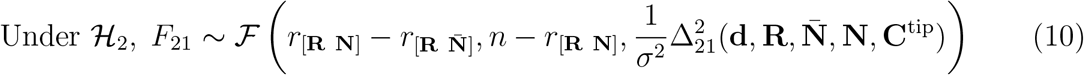

The noncentral coefficient are determined by ∆_10_ and ∆_21_, detailed in the Appendix. They depend on the network topology through the metric defined by **C**^tip^, and through the regression matrix **N**. Under the null hypothesis (*𝓗*_0_ for *F*_10_ and *𝓗*_1_ for *F*_21_), the statistics follow a central F distribution, and these ∆ terms are zero.

Because we know the exact distribution of our F statistics under the alternative hypothesis, we do not need to resort to simulations to assess the power of these tests. In the following, we present a theoretical power study.

#### Test 𝓗_0_ vs 𝓗_1_

We first studied the theoretical power to detect a single transgressive evolution effect, depending on the size *b* of this effect, and on the position of the hybridization event on the network. We considered 4 network topologies, using the same backbone tree than in the simulation study above, but adding only one hybridization event, occurring at various depths, from a very recent event affecting a single taxon to a very ancient event affecting 8 taxa (Fig. 5, top). The inheritance probability of this added hybrid branch was fixed to *γ* = 0.4. This *γ* parameter proved to have little influence to detect transgressive evolution (data not shown), for all the values tested, between 0 and 0.5. The underlying BM process had fixed ancestral value *µ* = 0, and variance rate *σ*^2^ = 1. Finally, for each network topology, we varied the transgressive evolution effect from 0 to 4, and computed the power of the test *𝓗*_0_ vs *𝓗*_1_ for three fixed standard levels (*α* in {0.01, 0.05, 0.1}). The range of effects (0 to 4) was chosen so that the power reaches 1 within this range for all 4 networks. This range is quite wide, compared to what could be considered a “biologically reasonable” effect size. As a comparison, we added a dashed line at *b* = 0.8 (Fig. 5), a value typically considered as being a “large” effect size (Cohen 1988). We can see that the power at *b* = 0.8 is rather small, hardly reaching 0.5 in the most favorable scenario. This reflects imbalance in group sizes, and power degradation due to phylogenetic correlation when reticulation is ancient (see Fig. 8 in the Appendix for a quantitative comparison). To give another benchmark, if the trait is measured on the log-scale, then *b* = log(2) *≈* 0.7 corresponds to a trait doubling because of transgressive evolution. We hence recommend doing a power study before collecting comparative data or after data collection, to determine which transgressive effects would likely go undetected due to a lack of power. We show in the next section how this can be done on a biological example, along with the empirical power observed. We also refer to the online supplementary material for practical ways to conduct a power analysis.

**Figure 5.**
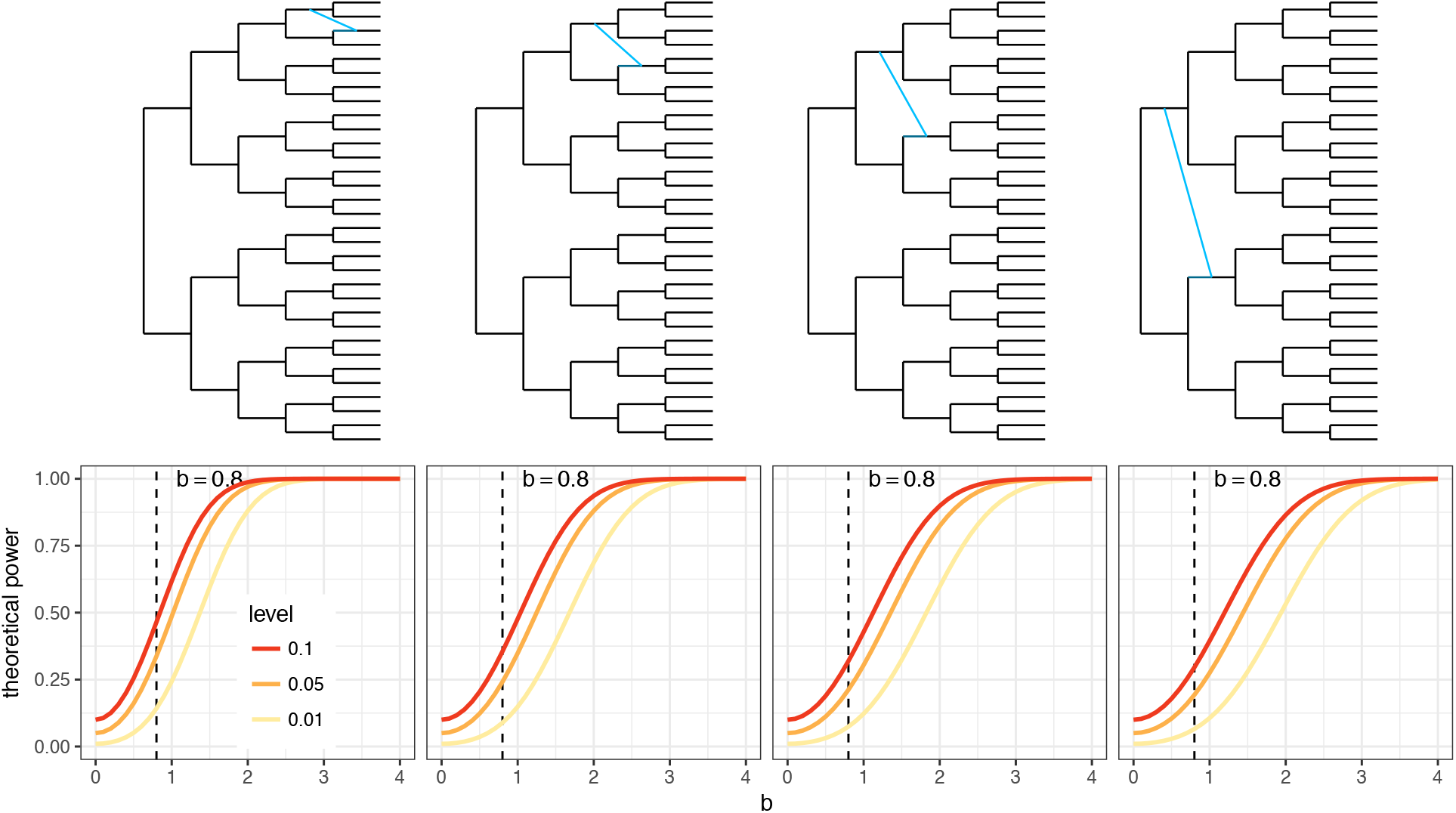
Theoretical power of the shared transgressive evolution test *𝓗*_0_ vs *𝓗*_1_, for four different networks topologies with inheritance probability *γ* = 0.4 (top), and a BM with ancestral value *µ* = 0 and variance rate *σ*^2^ = 1. The power of the test increases with the transgressive evolution effect *b* (bottom).

As expected, the power improves with the size of the effect, reaching approximately 1 for *b* = 4 in all scenarios (Fig. 5, bottom). In addition, the transgressive evolution effect seems easier to detect for *recent* hybridization events, even if they affect fewer tips. One intuition for that is that ancient hybridization effects are “diluted” by the variance of the BM, and are hence harder to detect, even if they affect more tips. This may be similar to the difficulty of detecting ancient hybridization compared to recent hybridizations.

#### Test 𝓗_1_ vs 𝓗_2_

We used a similar framework to study the power of the test to detect heterogeneity in the transgressive evolution effects. We used here the first 3 networks from the simulation study, with 32 tips and 2 to 8 hybridization events (Fig. 6, top), but with inheritance probabilities fixed to *γ* = 0.4. Transgressive evolution effects were set to **d** = *d***d**^*u*^, with **d**^*u*^ fixed to 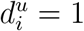 for *i ≤ h/*2 and 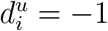 for *i > h/*2, *h* being the number of hybrids, which was even in all the scenarios we considered. Note that the average transgressive evolution effect was 0, because the 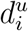 values sum up to 0. This allowed us to reduce the “strength of heterogeneity” to a single parameter *d*, which we varied between 0 and 4 (see appendices for the reduced expression of the noncentral coefficient). Like before, we computed the power of the test *𝓗*_1_ vs *𝓗*_2_ for three fixed standard levels (*α* in {0.01, 0.05, 0.1}).

**Figure 6.**
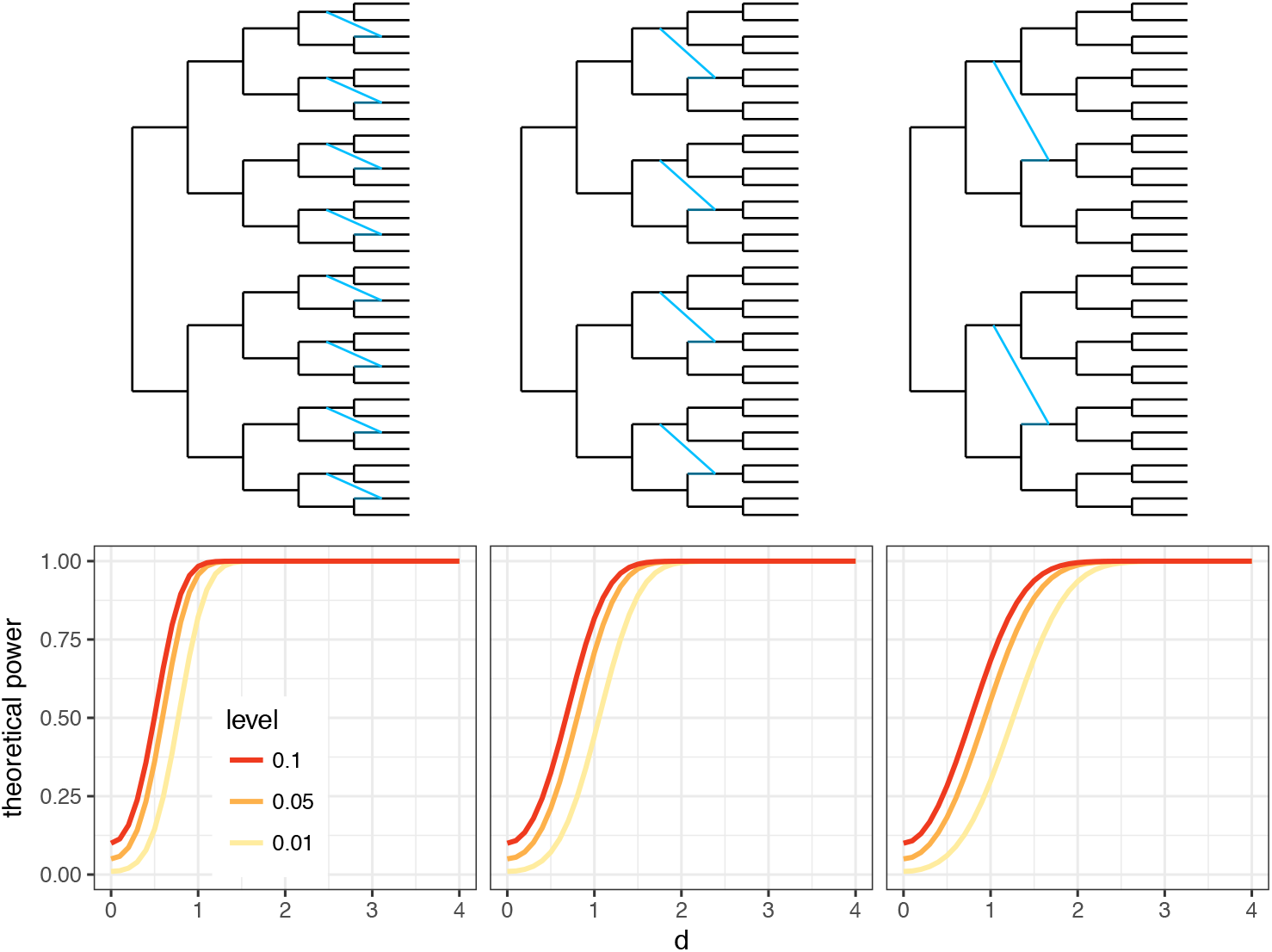
Theoretical power of the test for heterogeneous transgressive evolution *𝓗*_1_ vs *𝓗*_2_, for three different networks topologies with inheritance probability *γ* = 0.4 (top), and a BM with ancestral value *µ* = 0 and variance *σ*^2^ = 1. The power of the test increases with the heterogeneity coefficient *d*(bottom).

Figure 6 (bottom) shows a similar pattern: the test is more powerful for a high heterogeneity coefficient, and for recent hybridization events. For variation of about 2 in transgressive evolution, the power is close to one in all the scenarios considered here.

#### Power of hypothesis tests and confidence intervals

A major contribution of this work is to cast a network model of trait evolution in the well-studied framework of fixed-effects linear models, from which we borrow exact hypothesis tests and confidence intervals. Our power calculations provide insights to compare the information content across various networks, chosen to represent various possible hybridization scenarios. These calculations can be easily repeated on any phylogenetic network given a set of trait evolution parameters, as estimated from a data set for instance. For the analysis of a particular data set, we recommend the use of confidence intervals, which carry more information about the size of transgressive effects than the simple (non-)rejection of a hypothesis. These possibilities are illustrated in the next section.

## *Xiphophorus* fishes

### Methods

#### Network inference

We revisited the example in Solís-Lemus and Ané (2016) and re-analyzed transcriptome data from Cui et al. (2013) to reconstruct the evolutionary history of 23 swordtails and platyfishes (Xiphophorus: Poeciliidae). The original work included 24 taxa, but we eliminated *X. nezahualcoyotl*, because the individual sequenced in Cui et al. (2013) was found to be a lab hybrid not representative of the wild species *X. nezahualcoyotl* (personal communication). We re-analyzed their first set of 1183 transcripts, and BUCKy (Larget et al. 2010) was performed on each of the 8,855 4-taxon sets. The resulting quartet CFs were used in SNaQ (Solís-Lemus and Ané 2016), using *h* = 0 to *h* = 5 and 10 runs each. The network scores (negative log-pseudolikelihood) decreased very sharply from *h* = 0 to 1, strongly between *h* = 1 to 3, then decreased only slightly and somewhat linearly beyond *h* = 3 (Fig. 7, top left). Using a broken stick heuristic, one might suggest that *h* = 1 or perhaps *h* = 3 best fits the data. Given our focus on PCMs, we used both networks (*h* = 1 and 3) as well as the tree (*h* = 0) to study trait evolution, and to compare results across the different choices of reticulation numbers.

**Figure 7.**
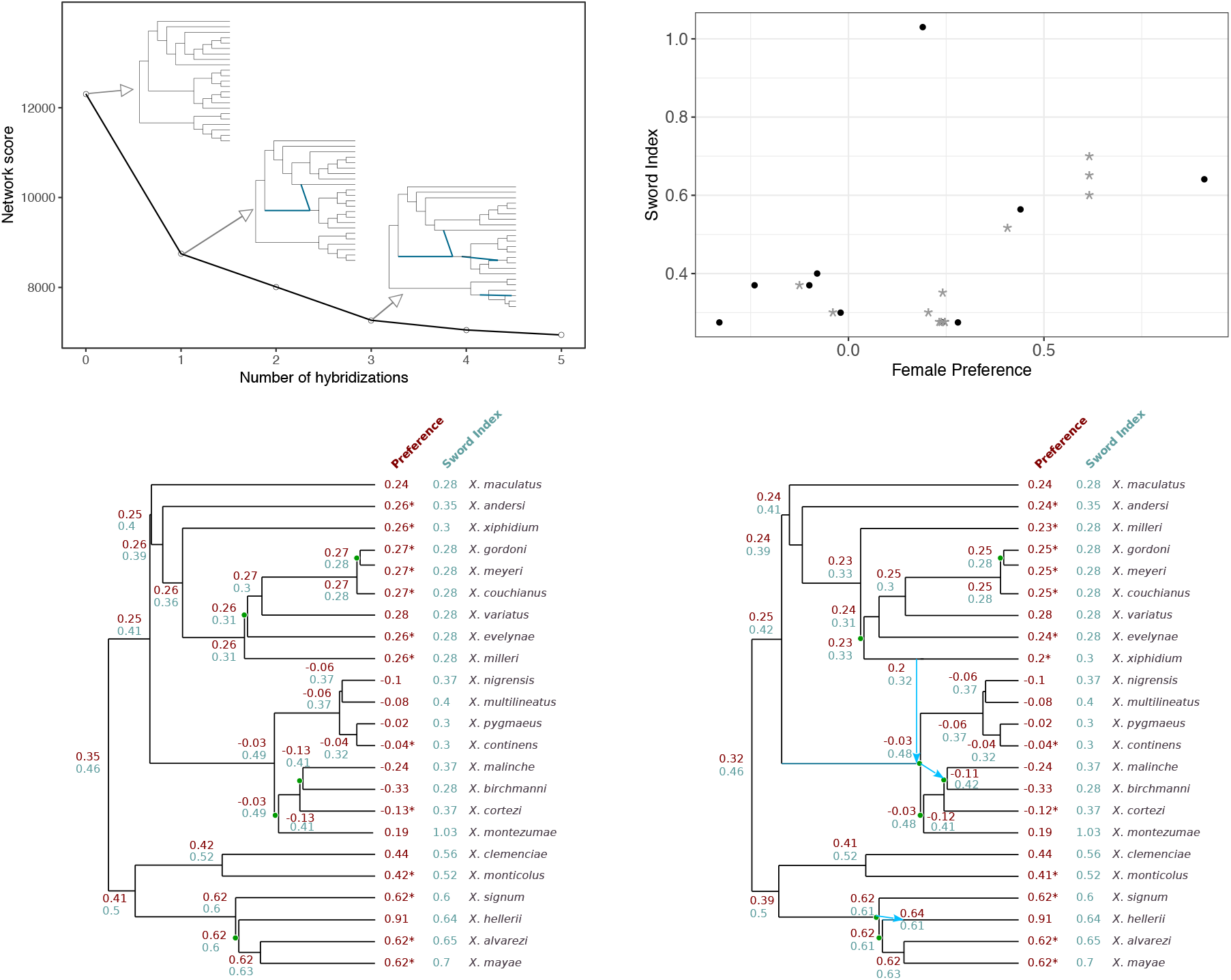
Results of the analysis on the fish dataset. Top left: negative pseudo log-likelihood score of the estimated networks with various numbers of hybridizations. Top right: scatter plot of sword index and female preference. Gray stars are taxa missing female preference data, for which female preference was predicted using ancestral state reconstruction of the trait on the network (independent of sword index). Bottom: ancestral state reconstruction of both traits, independently, using a BM model on the tree (*h* = 0, left) or on the network with *h* = 3 (right). Starred values indicate taxa with missing preference data, and imputed female preference values. Branches with an estimated length zero are indicated by a green dot, to show the network topologies.

#### Network calibration

SNaQ estimates branch lengths in coalescent units, which are not expected to be proportional to time, and are also not estimable for some edges (like external branches to taxa represented by a single individual). To calibrate the network, we estimated pairwise genetic distances between taxa, and then optimized node divergence times using a least-square criterion, as detailed below.

To estimate pairwise distances, individual gene trees were estimated with RAxML, using the HKY model and gamma-distributed rate variation among sites. For each locus, branch lengths were rescaled to a median of 1 to reduce rate variation across loci, before obtaining a pairwise distance matrix from each rescaled gene tree. Loci with one or more missing taxa were then excluded (leaving 1019 loci), and pairwise distance matrices were averaged across loci.

This average pairwise distance matrix was used to estimate node ages on each network (*h* = 0, 1, 3). The network pairwise distance between taxa *i* and *j* was taken as the weighted average distance between *i* and *j* on the trees displayed by the network, where the weight of a displayed tree is the product of the inheritance probabilities *γ_e_* for all edges *e* retained in the tree. We estimated node ages that minimized the ordinary least-squares mismatch between the genetic pairwise distances and the network pairwise distances.

#### Traits

With data presented in Cui et al. (2013) and following their study on sword evolution, we revisited the hypotheses that females have a preference for males with longer swords, and that the common ancestor of the genus *Xiphophorus* likely had a sword. Rather than using the methods of parsimony character mapping and independent contrasts as in Cui et al. (2013), we tested the effect of hybridization on the ancestral state reconstructions and the correlation between both traits using networks with zero, one or three hybridization events, using **phyloNetworklm**. For each network, the topology and branch lengths were assumed to be perfectly estimated, and fixed. We also tested for phylogenetic signal in both traits on all networks using Pagel’s *λ*, as well as for transgressive evolution, using the *F* statistics defined above. For the phylogenetic regression, more than half of the species were excluded because they lack information on female preference.

Along with the datasets used, two executables **IJulia notebooks** (.**ipynb**) files are provided in the online supplementary material (Dryad data repository doi:10.5061/dryad.60t0f), allowing the interested reader to reproduce all the analyses described here.

### Results

The *Xiphophorus* fish topologies with zero, one, and three hybridization events were calibrated using pairwise genetic distances (Fig. 7, bottom, for *h* = 0 and 3). With *h* = 1, the reticulation event did not necessarily imply the existence of unsampled or extinct taxa, so we constrained this reticulation to occur between contemporary populations (with an edge length of 0). For the network with *h* = 3, two reticulation events implied the existence of unsampled taxa, so we calibrated this network without constraint, to allow minor reticulation edges of positive lengths. Optimized branch lengths were similar between networks. Branch lengths were estimated to be 0 for some tree edges and some unconstrained hybrid edges, creating polytomies.

Using networks with 0, 1 or 3 hybridization events, we found a positive correlation between female preference and longer swords in males, but this relationship was not statistically significant (*h* = 0: *p* = 0.096; *h* = 1: *p* = 0.110; *h* = 3: *p* = 0.106). Ancestral state reconstruction of sword index shows the presence of a sword at the MRCA of each network because unsworded species were assigned a value of 0.275 in Cui et al. (2013) and the ancestral state in all networks was reconstructed to be 0.46. This reconstruction needs to be taken with caution, however, because 0.275 belongs in the 95% confidence interval for the ancestral sword index: [0.26, 0.66] for *h* = 3. This interval is wide when compared to the observed variation at the tips of the tree: [0.275, 1.03]. (Note that 0.275 is outside the 90% interval: [0.30, 0.63].) Phylogenetic signal was high for both traits with estimated *λ* = 1.0 on all networks (or above 1.0 with unconstrained maximum likelihood).

We also applied our tests for transgressive evolution on both traits, using the network with 3 hybridization events (Fig. 7, lower right). For the sword index, we found no evidence of transgressive evolution (*p* = 0.55 and *p* = 0.28, respectively, for homogeneous or heterogeneous transgressive evolution). This lack of evidence was reflected in the 95% confidence intervals for the transgressive shifts at the three hybridization events, which included 0: [−0.45, 0.06], [−0.20, 0.56] and [−0.34, 0.44]. However, we did find some evidence for an heterogeneous transgressive evolution effect for female preference. Testing *𝓗*_2_ against *𝓗*_1_ gives *p* = 0.0087. Testing *𝓗*_2_ against *𝓗*_0_ directly, we get *p* = 0.0064 (see the Appendix for a description of this third test, also based on a F statistic). However, transgressive evolution effects were in opposite directions (one positive and two negative), such that the common effect was not significantly different from 0: *𝓗*_1_ vs *𝓗*_0_ gave *p* = 0.11. Namely, the 95% confidence intervals for the shifts at the three hybridization events were [−0.57, −0.09], [−0.63, 0.10] and [0.12, 1.02]. Although these intervals are wide, the size of two of these effects is quite large: one negative and one positive by at least *∼* 10% of the observed variation at the tips ([−0.33, 0.91]). These large shifts match the fairly strong evidence for transgressive evolution from the F tests.

We computed the power of the tests (Fig. 5 and 6) but using the *Xiphophorous* network with three hybridizations, and using the estimated model parameters (including transgressive effects). The observed power for *H*_2_ vs *H*_0_ was low at 0.47 for the sword index but very high at almost 1.00 for the female preference.

## Discussion

### Impact of the Network

The results from the fish dataset analysis using a tree (*h* = 0) or a network (*h* = 1 or *h* = 3) show that taking the hybridization events into account has a small impact on the ancestral state reconstruction and on the estimation of parameters, both for the regression analysis and for the test for phylogenetic signal. This finding was corroborated by simulations: when we ignored hybridization events, using a tree while the true underlying model was a network, we found that the estimation of parameters *µ* and *σ*^2^ was only slightly affected (data not shown). These results may indicate that major previous findings, where a phylogenetic tree was used rather than a more appropriate network, are likely to be robust to a violation of the tree-like ancestry assumption. Our new model may simply refine previous estimates in many cases.

However, the structure of the network has a strong impact on the study of transgressive evolution. This is expected, as the model allows for shifts below each inferred hybrid. If one reticulation is undetected, or if one was incorrectly located on the network, then the model will be ill-fitted, leading to potentially misleading conclusions. As an example, we reproduced the analysis of transgressive evolution for female preference on the network with three hybridization events, but this time pruning the network, to keep only the taxa with a measured trait. Preference data were missing for species *X. signum*, *X. alvarezi* and *X. mayae*, such that *X. helleri* became the only species impacted by one of the reticulation event, which became a simple loop in the network. In other words, *X. helleri* was the only descendant of the reticulation, and also the closest relative of the hybrid’s parent among the remaining taxa. The reticulation could be dropped from the pruned network. This new and simplified network only retained the two hybridization events associated with negative shifts. As a consequence, and contrary to the conclusion we found in the main text, we found support for *homogeneous* transgressive evolution (*p* = 0.0071 for *𝓗*_1_ vs *𝓗*_0_), and no support for heterogeneous effects (*p* = 0.88 for *𝓗*_2_ vs *𝓗*_1_). This illustrates that caution is needed for the interpretation of tests of transgressive evolution, as those highly depend on the quality of the input network inference, which is a recognized hard problem.

### Network Calibration

To conduct PCMs, we developed a distance-based method to calibrate a network topology into a time-consistent network. This is a basic method that makes a molecular clock assumption on the input pairwise distance matrix. Important improvements could be made to account for rate variation across lineages, and to use calibration dates from fossil data, like in relaxed clock calibration methods for phylogenetic trees such as r8s (Sanderson 2003) or BEAST (Drummond et al. 2006). In our fish example, we averaged pairwise distances across loci, to mitigate a violation of the molecular clock that might be specific to each locus.

Our method estimated some branch lengths to be 0, thereby creating polytomies. This behavior is shared by other well-tested distance-based methods like Neighbor-Joining (Saitou and Nei 1987), which can also estimate 0 or even negative branch lengths.

We also noticed that several calibrations could fit the same matrix of genetic pairwise distances equally well, pointing to a lack of identifiability of some node ages. This issue occurred for the age of hybrid nodes and of their parent nodes. Branch lengths and node ages around reticulation points were also found to be non-identifiable by Pardi and Scornavacca (2015), when the input data consist of the full set of trees displayed by the network, and when these trees are calibrated. This information on gene trees can only identify the “unzipped” version of the network, where unzipping a reticulation means moving the hybrid point as close as possible to its child node (see Pardi and Scornavacca 2015, for a rigorous description of “canonical” networks). This unzipping operation creates a polytomy after the reticulation point. We observed such polytomies for two events in our calibrated network (Fig. 7, bottom right). Pardi and Scornavacca (2015) proved that the lack of identifiability is worse for time-consistent networks, which violates their “NELP” property (no equally-long paths). Lack of identifiable branch lengths around reticulations is thus observed from different sources of input data, and requires more study. Methods utilizing multiple sources of data might be able to resolve the issue. For instance, gene tree discordance is informative about branch lengths in coalescent units around reticulation nodes, and could rescue the lack of information from other input data like pairwise distances or calibrated displayed trees. More work is also needed to study the robustness of transgressive evolution tests to errors in estimated branch lengths.

### Comparison with Jhwueng and O’Meara (2018)

In their model, Jhwueng and O’Meara (2018) include hybridization events as *random* shifts. Using their notations, each hybrid *k* shifts by a coefficient log *β* + *δ_k_*, with *δ_k_* a random Gaussian with variance *ν_H_* : *δ_k_ ∼ N* (0, *ν_H_*). This formulation provides a *mixed effects* linear model, with shifts appearing as random effects. The effects of transgressive segregation, instead of being reflected in the mean as in our model, is then reflected in the extra variance *ν_H_* introduced after each hybrid. This extra term changes the structure of the variance matrix *C*, such that reticulation points do not necessarily induce a decrease in variance, like for the vanilla BM as shown in (2). In this framework, the test of heterogeneity (*𝓗*_2_ vs *𝓗*_1_) amounts to a test of null variance, *ν_H_* = 0. In the context of mixed effects linear models, such tests are also well studied, but are known to be more difficult than tests of fixed effects (Lehman 1986; Khuri et al. 1998). Assuming that the variance *ν_H_* is 0, our test for a common transgressive evolution effect (*𝓗*_1_ vs *𝓗*_0_) is then similar to the likelihood-based test for log *β* = 0 in Jhwueng and O’Meara (2018). A mixed-effect model is legitimate, although it might be more difficult to study theoretically, and its inference can be more tricky. Jhwueng and O’Meara (2018) indeed report some numerical problems, and rather large sampling error for both log *β* and *ν_H_*. Current state-of-the-art methods to infer phylogenetic networks cannot handle more than 30 taxa and no more than a handful of reticulation events (Hejase and Liu 2016). Hence, it might not be surprising that estimating a variance *ν_H_* for an event that is only observed two or three times is indeed difficult. On data sets with few reticulations, we believe that our fixed effect approach can be better suited. However, our approach adds a parameter for each hybridization event, whereas the random-effect approach of Jhwueng and O’Meara (2018) maintains only two parameters (mean and variance). As the available networks are likely to grow over the next few decades, this later approach might be preferable in the future.

### Comparison with pedigrees

There is an extensive literature for the analysis of phenotypic traits on individuals with a known pedigree (see Thompson 2000, and references therein). Pedigrees are highly detailed phylogenetic networks where nodes are individuals within a species. The ancestral state of a trait corresponds to the breeding value of a given ancestor. Our model is similar to the animal model for polygenic values. The correlation between the additive genetic (breeding) values of two individuals *i* and *j* was shown to be proportional to *A_ij_*, defined as twice the coefficient of kinship between *i* and *j* (Crow and Kimura 1970). This coefficient is the probability that two homologous genes picked at random from *i* and from *j* are identical by descent. The matrix **A** can be calculated recursively, taking individuals in the order in which they were born (preorder). Namely, if *i* has parents *a* and *b* then

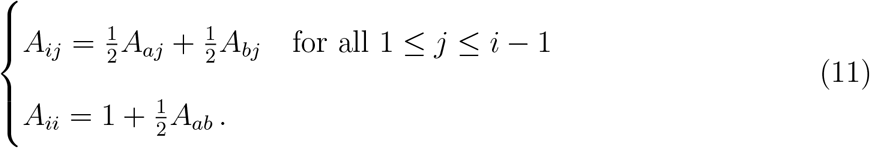

Next, 𝕧ar [**X**] = *σ*^2^**A** can be expressed as a linear recursive model: if individual *i* has parents *a* and *b*, then

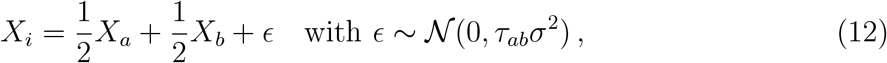

where *τ_ab_* = 1 − 0.25(*A_aa_* + *A_bb_*) (Henderson 1976; Mrode 2014, section 2.3). For a founder individual, *A_ii_* = 1 so *X_i_* is assumed to be normally distributed with variance *σ*^2^. In our framework, this model corresponds to Definition 1 on a network where each individual is a hybrid node, except for founders who act like roots with no parents. The network may have polytomies if an individual has multiple children. Each parent-child relationship is represented by a hybrid edge from parent *a* to child *i* with inheritance 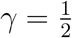 and length 2 *− A_aa_*. Since *A_aa_* depends on the pedigree of *a*, branch lengths in the network need to be computed recursively and cannot be specified a priori. The recursion (11) is equivalent to for covariance **C** in our model, given the specific *γ*s and branch lengths on the pedigree. The calculation of **A** was first derived by Wright (1922) using a path counting algorithm. We extend this algorithm to general networks in the Appendix, giving a path formula similar but different from (1).

The developments above show that the two main equations defining our model (the recursive and path methods for the variance computation) have a counterpart in the pedigree literature. However, there are important differences, both from a mathematical and a modeling point of view. Indeed, our model is more general than the pedigree model in that hybrid edges can have any inheritance *γ* not restricted to 1/2, tree edges can take any value to represent time ideally, and we can model transgressive evolution. In a pedigree network, branch lengths are such that the variance of all individuals is bounded by 2*σ*^2^. Non-inbred individuals have variance *σ*^2^, and inbred individuals have variance *σ*^2^*A_ii_* = *σ*^2^(1 + *f_i_*) depending on their inbreeding coefficient *f_i_*. On a general phylogenetic network, the BM variance grows indefinitely with time, a fact well recognized when using trees. This difference reflects their different biological justifications. The pedigree model was derived from a micro-evolutionary genetic mechanism within a population and one generation per edge, while the network model typically scales time in millions of years, and was developed from a heuristic model for macroevolution. Another major difference is data availability: trait data are typically observed at most nodes in a pedigree, but only at the tips of a phylogenetic network (with important computational consequences). Future work should build on the rich literature on pedigrees for faster computations on general networks (e.g. to invert **A**), or for expectation-maximization or Markov-chain Monte Carlo techniques.

### Extensions and perspectives

The BM model we presented here can be extended in many ways in order to account for various biological assumptions and mechanisms. First, keeping the vanilla BM, it could be interesting to look into other merging rules at reticulation points. For instance, instead of taking a weighted average, one could draw either one of the two parents’ trait for the hybrid, with probabilities defined by the weights *γ_a_* and *γ_b_* of the parents. If such a rule could be justified from a modelling point of view, further work would be needed to derive the induced distribution of the trait at the tips of the network.

Easy extensions could allow for rate variation. Following O’Meara et al. (2006) on trees, we could allow for rate variation across clades (or across separate parts of the network) by stretching or shrinking the edges in the same rate category by a common factor. One could then estimate the rate in each part of the phylogeny and then test if rates differ significantly. Extensions for rate variation over time could involve standard methods that rescale branch lengths, such as Pagel’s *κ* or *δ* as mentioned earlier. The early burst transformation (EB, Harmon et al. 2010) would be particularly valuable for studying adaptive radiation, to accommodate acceleration (or deceleration) of trait evolution (Blomberg et al. 2003), where the rate of evolution increases (or decreases) exponentially through time as 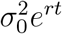, with *r <* 0 for early bursts followed by a slow down. Like Pagel’s *δ*, the EB model can be implemented via a transformation of node ages. A node of age *a* is given a new age of (*e^ra^ −* 1)*/r* under the EB model, so a branch of length *ℓ* starting at this node is rescaled to *e^ra^*(*e^ℓ^ −* 1)*/r*. Such transformations require a time-consistent network, in which the age of every node is well defined.

The Ornstein-Uhlenbeck (OU) process is popular to model trait evolution for the study of stabilizing selection, regime shifts, and convergent evolution (e.g. Hansen 1997; Butler and King 2004; Beaulieu et al. 2012; Khabbazian et al. 2016; Bastide et al. 2018). The OU process has extra parameters compared to the BM: a primary optimum *θ* representing an adaptive peak, and a rubber band parameter *α* that controls how fast the trait is pulled toward its optimum. Extending our network model to an OU process is complicated because the mean of the OU process, not just the variance, changes over time along each lineage. After evolving for time *ℓ*, the trait *X_b_* of the OU process has a mean that depends on both the ancestral value *X_a_* and the primary optimum: *e^−αℓ^ X_a_* + (1 *− e^−αℓ^*)*θ*. What trait value would be biologically realistic at reticulation points? For an OU with one single optimum *θ* over the whole tree, the ancestral trait at the root can be assumed to be centered on *θ*, such that the mean trait value is *θ* at all nodes. In this case, the weighted average merging rule could be adopted. But how should transgressive evolution be modeled? With the OU process, shifts have been traditionally considered on its parameters (like *θ*) rather that directly on the trait itself *X*, as we did for the BM (Butler and King 2004; Beaulieu et al. 2012). If a transgressive evolution shift is allowed on the optimum value, this would result in several optima on different regions of the network, which might not capture biological realism. A related problem is to find a realistic merging rule for reticulations between two species evolving in two different phylogenetic groups with different optima.

PCMs rely on two fundamental components: the species relationship model (tree or network), and the model of trait evolution. Here, we showed how a network could be used instead of a tree. Our study sets up a rigorous and flexible theoretical framework for PCMs on phylogenies with reticulations. Taking the simplest model for continuous trait evolution the BM with fixed variance – we showed how some standard tools, such as phylogenetic regression or test of phylogenetic signal, can be extended to take reticulation into account.

We also discussed issues that are specific to networks and offered new tools to deal with them, such as tests for transgressive evolution. The numerous improvements that have been developed for PCMs on trees should be adapted to phylogenetic networks, starting with support for measurement error or intra-specific variation (as in, e.g. Lynch 1991; Ives et al. 2007; Felsenstein 2008; Goolsby et al. 2017); multivariate processes (Felsenstein 1985; Bartoszek et al. 2012; Clavel et al. 2015) and developments mentioned above. Unexplored and more challenging questions will be to analyze geographical traits (biogeography) or to correlate trait evolution with diversification, when the phylogeny has reticulations. A salient point to be careful about will be the merging rule one might adopt for all these processes. Our work opens a door for much needed future work for trait evolution on phylogenetic networks.

## Acknowledgments

We thank Frank Burbrink and all participants of the spotlight session on the impact of gene flow and reticulation in phylogenetics, at the 2017 Evolution meeting. PB thanks Mahendra Mariadassou and St´ephane Robin for enlightening comments on an early version of this work, and Tristan Mary-Huard for useful discussions about the linear mixed model. The authors thank Mohammad Khabbazian for insights on the topological sorting algorithm, and Guilherme Rosa for discussions on the pedigree model. This manuscript also highly benefited from feedback by Frank Burbrink, Brian O’Meara and two anonymous reviewers.

## Funding

The visit of PB to the University of Wisconsin-Madison during the fall of 2015 was funded by a grant from the Franco-American Fulbright Commission. This work was funded in part by the National Science Foundation (DEB 1354793) and by a Vilas Associate award to CA from the University of Wisconsin-Madison.

## Supplementary Material

Supplementary material, including data files and two executables IJulia notebooks that reproduce the analyses conducted in the main text, can be found in the Dryad data repository at http://datadryad.org, doi:10.5061/dryad.nt2g6.

## Proof of the Variance Formula and Algorithm

We prove here both formula (1) for the BM variance matrix and Proposition 1 giving an efficient algorithm to calculate this matrix. We do so by induction on the number of nodes in the network: *N* = *n* + *m*. When the network is made of a single node *i* = 1, equation (1) and Proposition 1 are obviously correct. We now assume that these results are correct for any phylogenetic network with up to *N −* 1 nodes, and we consider a network with *N* nodes. When these nodes are sorted in preorder, the last node *i* = *N* is necessarily a tip (with no descendants), so removing it and its parent edges from the original network gives a valid phylogenetic network with *N −* 1 nodes. Using the same notations as in the main text, we can focus on the case *i* = *N*. Because of the preorder, there is no directed path from *i* to *j* for any *j* < *i*. We use here the formulas of Definition 1, and assume *σ*^2^ = 1 without loss of generality.

- If *i* is a hybrid node, then *X_i_* = (*γ_e_a__ X_a_* + *γ_e_b__ X_b_*) + (*γ_e_a__ ∈_a_* + *γ_e_b__ ∈_b_*), with *∈_k_ ∼ 𝒩* (0, *ℓ_ek_*), and *∈_k_* independent of the all values *X_j_* in the subnetwork (*j* < *i*) for *k* = *a* and *k* = *b*. Because of the preorder, *a* < *i* and *b* < *i*. Then:

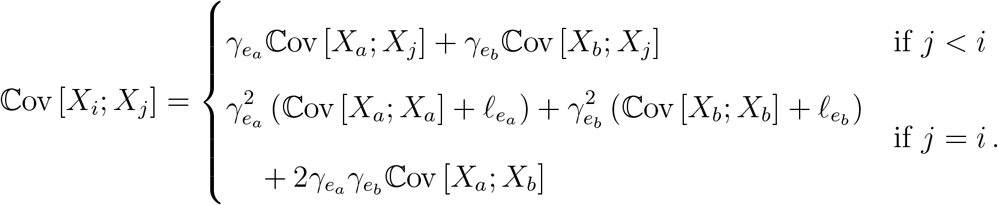

This proves (4) in Proposition 1. Next, we focus on proving (1). Note that it is valid by induction for all nodes in the subnetwork, and we just need to prove it for *i* = *N* and any *j* ≤ *i*. By induction, (1) holds for *a*, *b*, and any *j* < *i*. Then, because *a* and *b* are the only parents of *i*, any path *p_i_* from the root to *i* must go through *a* and *e_a_*, or through *b* and *e_b_* (and not both). In other words:

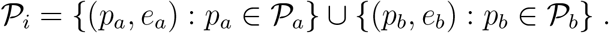

Now considering node *j* < *i* and a path *p_j_* from the root to *j*, *p_j_* cannot go through *i* so it cannot go through *e_a_* or *e_b_*. Therefore, the shared edges between *p_j_* and *p_i_* = (*p_a_, e_a_*) are exactly the same edges as those shared between *p_j_* and *p_a_*, and the shared edges between *p_j_* and *p_i_* = (*p_b_, e_b_*) are also the same as those shared between *p_j_* and *p_b_*. For *j* < *i*, we get:

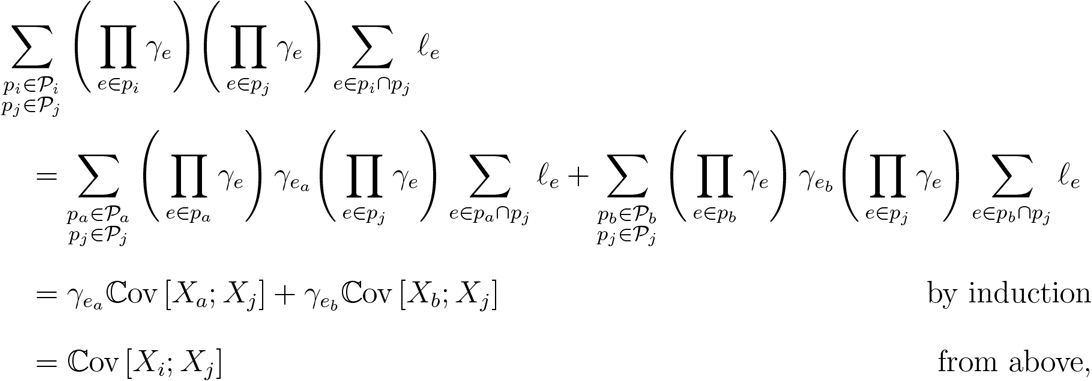

proving (1) for *i* = *N* and *j* < *i*. For *j* = *i* = *N*, we similarly decompose the set of paths *𝒫_i_* into two sets, either going through *a* or through *b*:

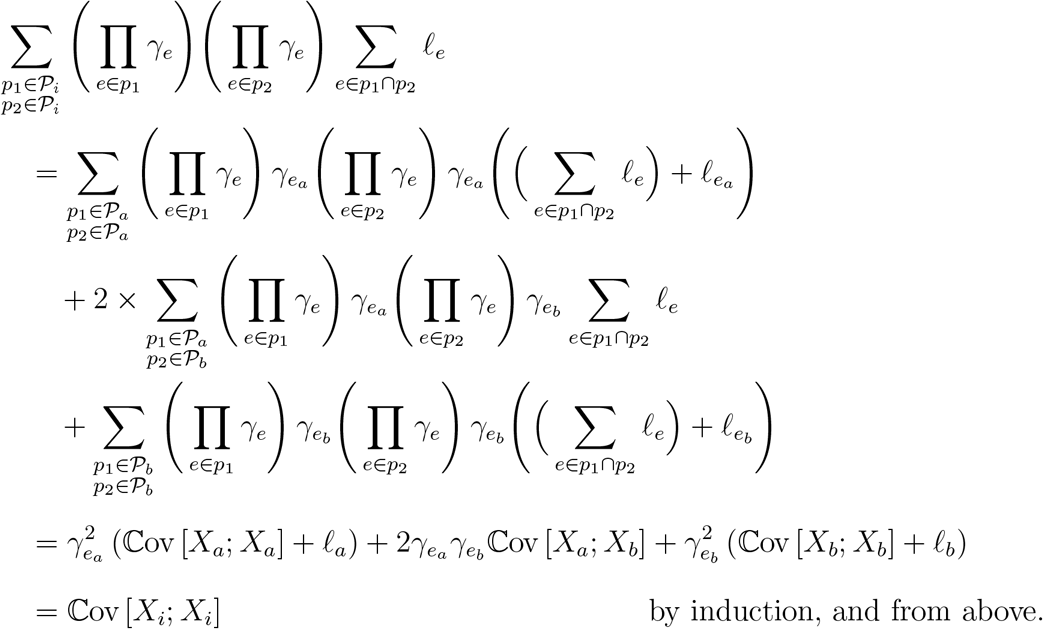

Where we used the equality 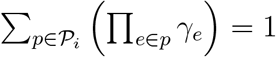 to go from the first to the second line. This completes the proof of (1), for *i* = *j*, and for the last case when *i* is a hybrid node.

- If *i* is a tree node, then *X_i_* = *X_a_* + *∈_a_*, with *∈_a_ ∼ 𝒩* (0, *ℓ_e_a__*), *∈_a_* independent of the values *X_j_* in the subnetwork (*j < i*). A tree node can be seen as a particular case of a hybrid node by taking *γ_e_a__* = 1, and creating an imaginary edge *e_b_* from *b* = *a* to *i* parallel to *e_a_* (for instance), with *γ_e_b__* = 0. This allows us to write *X_i_* = (*γ_e_a__ X_a_* + *γ_e_b__ X_b_*) + (*γ_e_a__ ∈_a_* + *γ_e_b__ ∈_b_*) as before. Equation (3) in Proposition 1 then follows directly as a limit case. Equation (1) also follows from the same derivation as for a hybrid node, because all the paths going through the new edge *e_b_* contribute nothing due to *γ_e_b__* = 0.

## Variance Reduction

Here, we prove Formula (2). As in the main text, consider a time-consistent network. For tip *i*, let *t_i_* be the length of any path from the root to *i*. If the history of tip *i* involves one or more reticulations then take any two paths *p_i_* and *q_i_* in *P_i_*. We have: ∑_*e*∈*pi*∩*qi*_ ℓ_*e*_ ≤ ∑_*e*∈*pi*_ ℓ_*e*_ = *t_i_*, with a strict inequality if and only if *p_i_* and *q_i_* are different paths. Seeing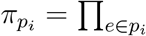 *γ_e_* as the probability associated with the path *p_i_* (with ∑_*pi*∈𝒫*i*_ π_*pi*_ = 1), we get from Equation (1):

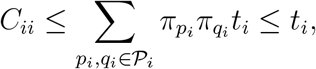

with the equality fulfilled if and only if there is a unique path from the root to taxon *i*, i.e. if *i* has no hybrid ancestry.

## Pagel’s *λ* Variance

*Proof of Proposition 2*. In Equation 1, the first equation is straightforward, because all the edges shared by the paths to *i* and to *j* are internal edges, whose lengths are multiplied by λ. Now take a tip node *i*. The first step of the transformation ensures that *i* is a tree node. Let *a* be its parent node, and parent branch *e_a_*. From the recursive formula given in Proposition 1, the variance at node *i* is proportional to:

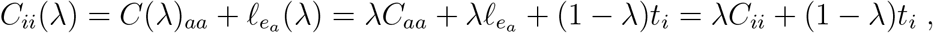

hence the announced formulas.

## Shifted BM model with the Descendence Matrix

*Proof of Formula* (7). The shifts are fixed, so they do not impact the variance structure of the traits, and we only need to show that 𝔼[Y] = **T∆**. Here, we prove a slightly more general formula on the complete vector of trait values at all the nodes, that is: 𝔼[X] = **U∆**. The original equality is easily derived from this one by keeping the tip values only.

We show this equality recursively. Assume that the nodes are numbered in preorder. Denote by **U**^*i*^ the *i^th^* row-vector of **U**. Node *i* = 1 is the root, which is the descendant of no other node than itself, so

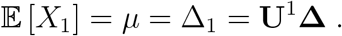

We now assume that E[X_*j*_] = **U**^*j*^ **∆** for all nodes *j < i*, and we seek to prove that this property is also true for node *i*.

- If *i* is a tree node, then denote by *a* its unique parent and by *e_a_* the edge from *a* to *i*. For any node *k ≠ i*, *P_k→i_* = {(*p_a_, e_a_*) : *p_a_ ∈ P_k→a_*}. Since *e_a_* is a tree edge with *γ_e_a__* = 1, we get from definition 3 that:

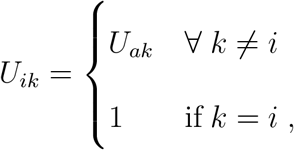

hence

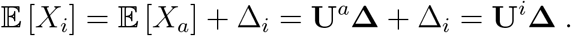

- If *i* is a hybrid, then denote by *a* and *b* its two parents, by *e_a_* and *e_b_* the corresponding edges, with coefficients *γ_e_a__* and *γ_e_b__*. Then for any node *k ≠ i*, we have: *P_k→i_* = {(*p_a_, e_a_*) : *p_a_ ∈ P_k→a_*} ∪ {(*p_b_, e_b_*) : *p_b_ ∈ P_k→b_*}, and using definition 3:

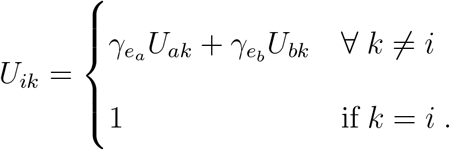 Since no shift can occur on the hybrid branches, ∆_*i*_ = 0 by convention and:

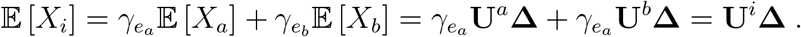 This ends the recursion, and the proof of (7).

Note that this proof also gives an efficient recursive way to compute the descendence matrix **U**.

### Mixed model formulation

The transgressive evolution model (7) and the phylogenetic linear regression (5) have a residual term with variance **C**^tip^ structured by the network. In the same way as phylogenetic regression on trees (Lynch 1991), (5) can be seen as a linear mixed model. It is straightforward to prove by induction that model (7) written for the entire network is equivalent to

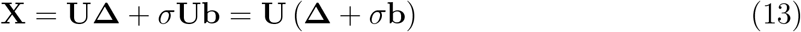

where **b** is a random vector with *independent* entries *b_i_ ∼ 𝒩* (0, *L_i_*); and *L_i_* = *ℓ_e_* if *i* is a tree node with parent branch *e*, 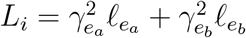 if *i* is a hybrid node with parent branches *e_a_* and *e_b_*, and *L_ρ_* is 0 if the root *ρ* has a fixed trait value, and 1 if the root is taken random with variance *σ*^2^. Besides giving an alternative statistical form to (7) (and more generally), this reformulation allows us to prove an alternative path formula for the variance matrix:

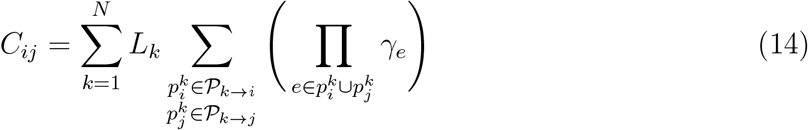

where *𝒫_k→ i_* is as in Definition 3, and where we chose by convention that 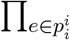 γ_e_ = 1. To*i* prove (14), we use (13) to write 𝕧ar [**X**] = *σ*^2^**ULU**^*T*^ where **L** is the diagonal matrix with entries the *L_i_* defined above. Hence, for any nodes *i* and *j*:

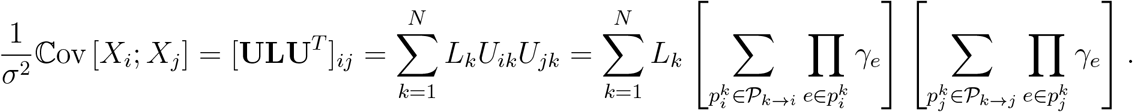

Note that this formulation has both fixed and random effects, but lacks an additional residual error term **E** (with independent entries) to match the phylogenetic mixed model (Lynch 1991). This error term could represent other sources of variation, such as intra-specific variation or measurement error. Including this term was proven to be crucial in PCM analyses on trees (Silvestro et al. 2015), and including it on networks should be the focus of future work.

## F Test for Transgressive Evolution

The F statistics used in Section Transgressive Evolution have the following expression:

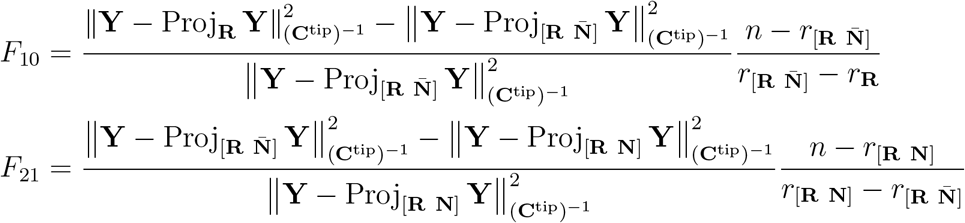

where Proj_**M**_ denotes the projection onto the linear space spanned by the columns of matrix **M**, with respect to the metric defined by 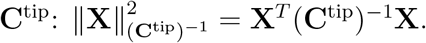 In other words, for any vector **X**:

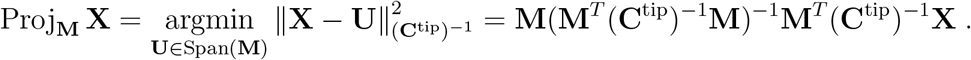

These statistics follow a noncentral F distribution as given in (9) and (10) of the main text, where

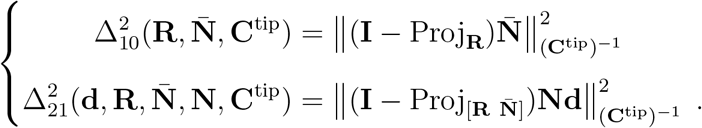

When studying the power of the test *𝓗*_1_ vs *𝓗*_2_, we took **d** = *d***d**^*u*^, so that the noncentral coefficient is:

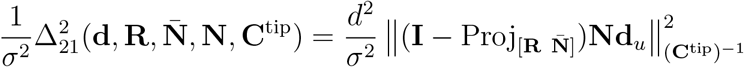

and, as the networks are fixed, it only varies with the heterogeneity coefficient *d*.

Note that a third statistic, *F*_20_, can be defined in a similar way to test *H*_2_ vs *H*_0_ directly. We first re-write the linear model as:

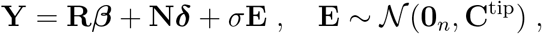

where there are no constraints on coefficients ***δ***. Then the *F* statistic can be written as:

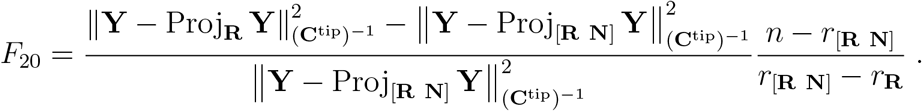

In the same way, it follows under *𝓗*_2_ a noncentral F distribution:

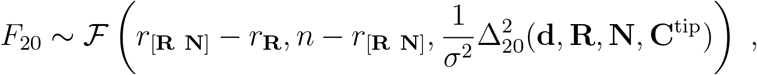

with

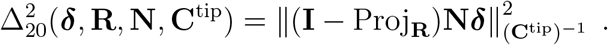

Thanks to the flexible framework provided by the GLM ftest function, all these tests are readily implemented, as long as one can fit the three models (*𝓗*_0_, *𝓗*_1_, and *𝓗*_2_).

### Comparison with independent species

We compared the power of our test for phylogenetically correlated species, to a situation where all the species would be independent. With independent units and for parameters in Figure 5, we get 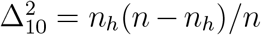, where *n* = 32 is the total number of species, and *n_h_ ∈* {1, 2, 4, 8} is the number of species with a hybrid ancestry. The effect size *b* is the mean difference between species with a hybrid ancestry, and species with no hybrid ancestry, assuming variance *σ*^2^ = 1 within groups. This allows us to compute the power to detect a shift (group difference) at various values of *b*. This power can then be compared to that obtained under phylogenetic correlation. The network structure tends to degrade the power when there are more than 4 species impacted by the shift (Fig. 8). Even for independent species, the small group sizes make for generally quite low power. In the most favorable situation, the power to detect an effect of size *b* = 0.8 is only 0.47 (Fig. 8, right-most panel, dashed curve). In the more standard situation where the two groups each have 30 individuals, an effect of size *b* = 0.8 would be detected with power 0.86, which would typically be considered as sufficient. Interestingly, in the two networks on the left where only one or two species are impacted by transgressive evolution, the network structure actually improves the power. In these networks, the species with hybrid ancestry have very close sister clades, which provide information about the ancestral trait just before the transgressive shift. The high correlation between the recent hybrid and its sisters improves the power to detect the shift.

**Figure 8:**
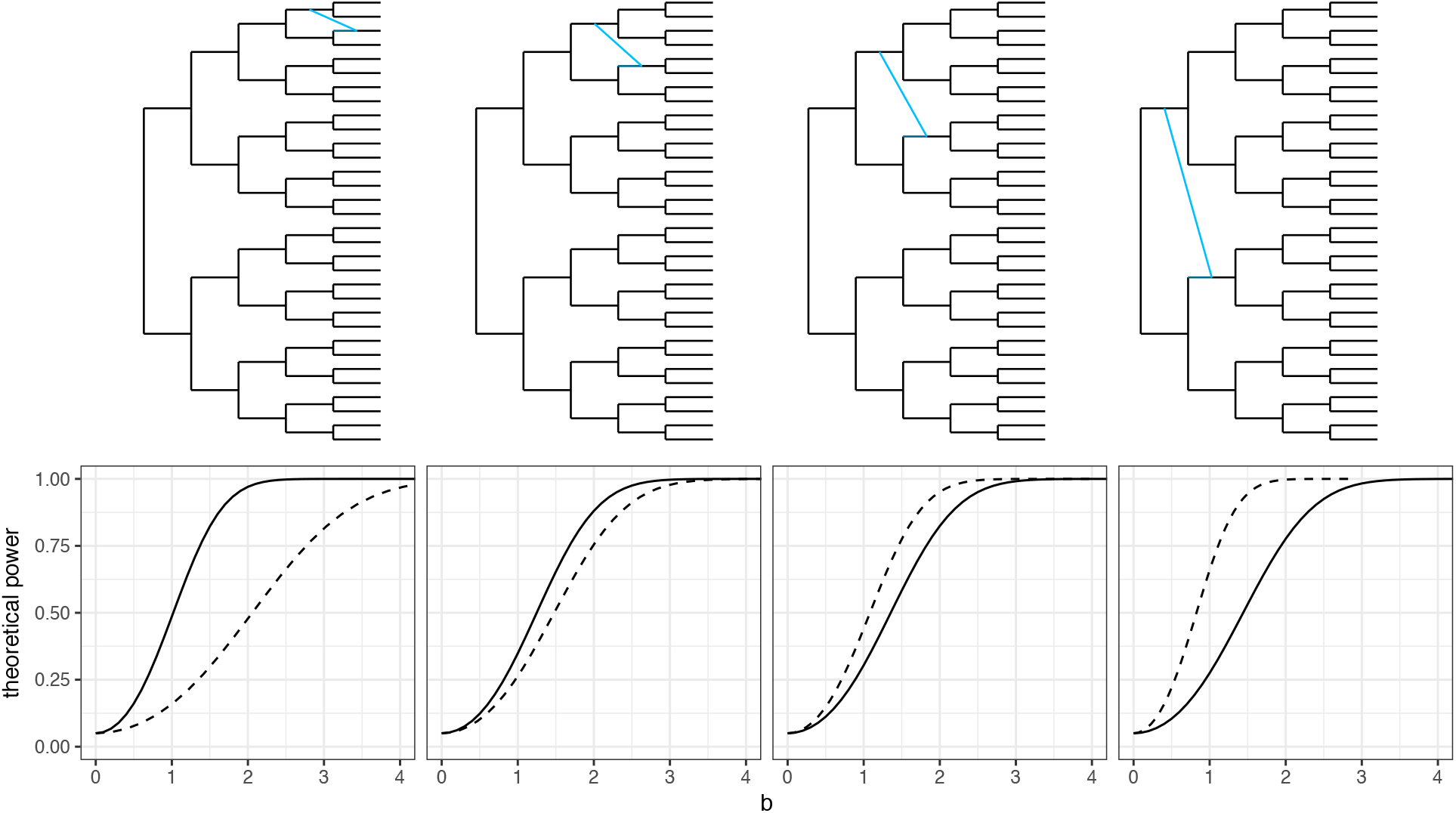
Theoretical power of the shared transgressive evolution test *H*_0_ vs *H*_1_, as in Figure 5, for a level of 0.05. The solid curve shows the actual test on the network. The dashed curve shows the power if the species were all independent, and if the traditional F test (or Student T test) were used to detect a shift affecting the species with hybrid ancestry. Under independence, the power is highly dependent on sample sizes, with more balance providing higher power. The network structure degrades the power compared to the independent case, except when the hybridization is recent, in which case the dependence structure helps. Under phylogenetic correlation, power is affected by the age of the hybridization and by the imbalance in group sizes (with opposing effects here).

## Link with pedigrees

We first note that pedigrees may contain individuals with a single known parent, a case omitted from the main text for conciseness. This case is easily modeled in the network by including a node for the unknown parent, considering it as a founder individual. We also note that in many reference publications, (12) is stated with *τ_ab_* simplified to 1*/*2 (Thompson 2000; Thompson and Shaw 1990, e.g.). This simplified model is correct only if the pedigree contains no inbred individuals.

### Path formula

We present here a path formula that is analogous to the path counting method on pedigrees from Wright (1922) (equation (3.1) in Thompson 2000), generalized to phylogenetic networks. For any *i ≠ j*, we show below that:

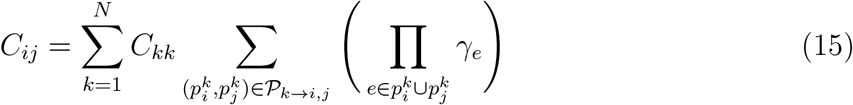

where *P_k→i,j_* is the set of pairs of directed paths 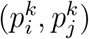such that 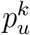goes from *k* to *u* and such that 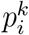 and 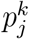 are disjoint, in the sense that they do not share any node other than *k* (hence they do not share any edge). When applied to a pedigree, the last term 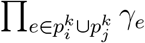simplifies to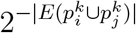where *|E*(*p*)*|* is the number of edges in path *p*. Unlike (1) and (14), says nothing about *C_ii_*. Like (14), (15) is applicable to networks with multiple roots.

*Proof*. We prove (15) by induction. For a network with a single node, there is nothing to prove. For a network with *N* nodes, we preorder the nodes. By induction, (15) holds for the subnetwork made of nodes {1, …, *N* − 1}. Next, we need to prove that (15) holds for any *i < N* and for *j* = *N*.

- If *N* is a tree node with parent *a*, then *C_iN_* = *C_ia_* for any *i ≠ N*, from (3). If, further, *i ≠ a* then

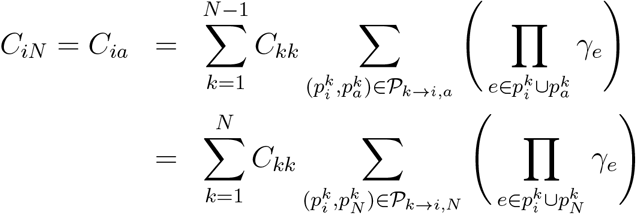

thus proving (15). The last line comes from the fact that any path 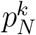 is the union of one path 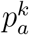 with the edge connecting *a* to *N*, which has *γ* = 1, and 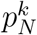 is disjoint with 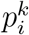 if and only if 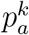 is disjoint with 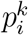. Also, the contribution of node *k* = *N* is 0 because the set of paths from *k* = *N* to *i* is empty. If *i* = *a*, then (15) holds again because it simplifies to *C_aa_*: the only node *k* with a non-empty pair *P_k→a,N_* is *k* = *a*, for which there is a single pair 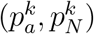 where 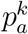 has node *a* only (no edges), and 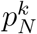has *a* and *N* (and a single edge). For this pair, the contribution is *γ* = 1.
- If *N* is a hybrid node with parents *a* and *b*, then *C_iN_* = *γ_a_C_ia_* + *γ_b_C_ib_* for *i ≠ N* from (4). Further, if *i* ≠ *a* and *i ≠ b*, then we can apply (15) to (*i, a*) and (*i, b*) to get:

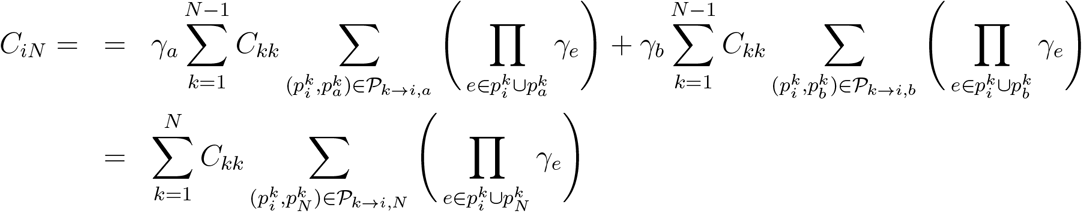

thus proving (15). The last line comes from the fact that any path 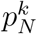 is the union of one path 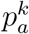 with the edge connecting *a* to *N*, which has *γ* = γ_a_, or the union of one path 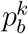 with the edge connecting *b* to *N*, which has *γ* = γ_b_. Also, 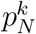 is disjoint with 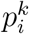 if and only if 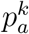 (resp. 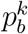) is disjoint with 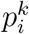. Also, like before, the contribution of *k* = *N* is 0 because the set of paths from *k* = *N* to *i* is empty. Next, if *i* = *a*, then (15) holds again:

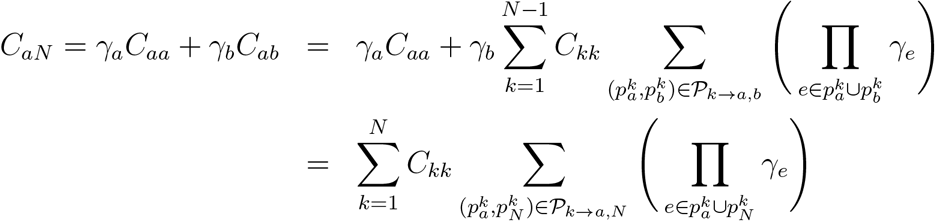

because if 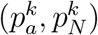 is in *P_k→a,N_* and if *k ≠a*, then 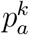 is required to be disjoint from 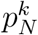, so 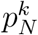 must be the union of a path 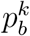 with the edge from *b* to *N* (with *γ* = *γ*_b_), and 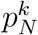 is disjoint from 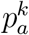 exactly if 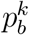 is disjoint from 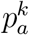. Also, the contribution of *k* = *a* is 0 in *C_ab_*, and γ_a_C_aa_ on the last line. The argument for *i* = *b* is analogous to the case *i* = *a*.

